# Distinct neural circuits establish the same chemosensory behavior in *C. elegans*

**DOI:** 10.1101/2021.08.17.456617

**Authors:** Navonil Banerjee, Pei-Yin Shih, Elisa J. Rojas Palato, Paul W. Sternberg, Elissa A. Hallem

**Affiliations:** Department of Microbiology, Immunology, and Molecular Genetics, University of California, Los Angeles, CA 90095; Molecular Biology Institute, University of California, Los Angeles, CA 90095; Division of Biology and Biological Engineering, California Institute of Technology, Pasadena, CA 91125; Department of Ecology, Evolution and Environmental Biology and Zuckerman Mind, Brain, Behavior Institute, Columbia University, New York, NY 10027

## Abstract

Animals frequently exhibit the same behavior under different environmental or physiological conditions. To what extent these behaviors are generated by similar vs. distinct mechanisms is unclear. Moreover, the circumstances under which divergent neural mechanisms establish the same behavior, and the molecular signals that regulate the same behavior across conditions, are poorly understood. We show that in *C. elegans*, distinct neural mechanisms mediate the same chemosensory behavior at two different life stages. Both dauer larvae and starved adults are attracted to carbon dioxide (CO_2_), but CO_2_ attraction is mediated by distinct sets of interneurons at the two life stages. Some interneurons mediate CO_2_ response only in dauers, some show CO_2_-evoked activity in adults and dauers but contribute to CO_2_ response only in adults, and some show CO_2_-evoked activity that opposes CO_2_ attraction in adults but promotes CO_2_ attraction in dauers. We also identify a novel role for insulin signaling in establishing life-stage-specific CO_2_ responses by modulating interneuron activity. Further, we show that a combinatorial code of both shared and life-stage-specific molecular signals regulate CO_2_ attraction. Our results identify a mechanism by which the same chemosensory behavior can be generated by distinct neural circuits, revealing an unexpected complexity to chemosensory processing.

## Introduction

Many animals engage in the same behavior under widely varying environmental and physiological contexts. For example, juveniles and adults of the same species often engage in similar foraging or escape behaviors despite their vastly different physiological states (Pereira and Sokolowski 1993, Jackson and MacMillan 2000, Bradley et al. 2013, Stern et al. 2017). When an animal generates the same behavior under different circumstances, it is often assumed that the behavior is established by the same neural mechanisms regardless of context. Although electrophysiological studies have demonstrated that distinct circuit mechanisms can generate similar network outputs (Prinz et al. 2004, Bucher et al. 2005, Saideman et al. 2007, Goaillard et al. 2009, Rodriguez et al. 2013, Hamood and Marder 2014, Marder et al. 2015, Cropper et al. 2016, Wang et al. 2019), the impact of such mechanisms on behavior is incompletely understood. Prior studies conducted in the free-living nematode *Caenorhabditis elegans* have shown that, in some instances, the same behaviors may arise from different neural mechanisms (Beverly et al. 2011, Trojanowski et al. 2014). In humans, children and adults were found to use different brain areas to perform the same word generation task (Brown et al. 2005), indicating that distinct neural pathways driving the same behavior may be widespread throughout the animal kingdom. The most common mechanism by which different circuits can give rise to the same behavior involves degeneracy, where multiple circuit components are capable of driving the same circuit output (Beverly et al. 2011, Trojanowski et al. 2014, Cropper et al. 2016). However, whether other mechanisms, such as the repurposing of circuit components, can be used to generate the same behavior across contexts remains unclear. In addition, the specific conditions under which such mechanisms are likely to be revealed are poorly understood, and the molecular mechanisms that enable distinct neural circuits to drive the same behavior across conditions have not been investigated.

To address this question, we used the behavioral responses of *C. elegans* to carbon dioxide (CO_2_) as a model system. *C. elegans* has a small nervous system with a well-characterized connectome (White et al. 1986, Varshney et al. 2011, Cook et al. 2019). In addition, *C. elegans* responds robustly to a diverse array of sensory cues, including CO_2_ (Banerjee and Hallem 2020). CO_2_ is an ambiguous chemosensory cue for *C. elegans*, as elevated CO_2_ levels in its natural habitat may signal food, predators, pathogens, or conspecifics (Carrillo and Hallem 2015, Banerjee and Hallem 2020). Accordingly, *C. elegans* shows flexible responses to CO_2_ such that CO_2_ can be either attractive or repulsive depending on immediate context, prior experience, and life stage (Bretscher et al. 2008, Hallem and Sternberg 2008, Hallem et al. 2011, Carrillo et al. 2013, Kodama-Namba et al. 2013, Guillermin et al. 2017, Rengarajan et al. 2019). For example, well-fed *C. elegans* adults are repelled by CO_2_, while starved adults are attracted to CO_2_; this shift in CO_2_ response valence occurs over the course of a few hours following food deprivation (Bretscher et al. 2008, Hallem and Sternberg 2008, Rengarajan et al. 2019).

Under adverse environmental conditions such as absence of food, high temperature, and overcrowding, *C. elegans* enters the developmentally arrested dauer larval stage (Golden and Riddle 1984, Hu 2007). Dauer entry is accompanied by a dramatic reprogramming of internal physiology that promotes developmental arrest and prolonged survival under unfavorable conditions (Fielenbach and Antebi 2008). Like starved adults, dauer larvae are robustly attracted to CO_2_ (Hallem et al. 2011). In adults, both CO_2_ attraction and CO_2_ avoidance are mediated by the same neural circuit, and CO_2_ response valence is determined by experience-dependent modulation of interneuron activity (Guillermin et al. 2017, Rengarajan et al. 2019). However, the neural circuit that mediates CO_2_ attraction in dauers had not been investigated.

Here, we show that dauers and starved adults use distinct neural circuits to generate attractive behavioral responses to CO_2_. While both dauers and adults require the CO_2_-detecting BAG sensory neurons for CO_2_ attraction, they require distinct sets of downstream interneurons. These interneurons show life-stage-specific differences in their functional properties as well as life-stage-specific roles in regulating CO_2_-evoked behavior. In particular, the AIB and AVE interneurons exhibit dauer-specific CO_2_-evoked activity and promote CO_2_ attraction selectively in dauers. In contrast, the CO_2_-evoked activity of the RIG interneurons opposes CO_2_ attraction in adults but promotes CO_2_ attraction in dauers, while the AIY interneurons show CO_2_-evoked activity in both dauers and adults but are required for behavior specifically in adults. Differences in the functional composition of the CO_2_ circuit in dauers vs. adults arise in part from the formation of a dauer-specific electrical synapse (Bhattacharya et al. 2019). In addition, we elucidate a novel role for insulin signaling, mediated by the insulin receptor homolog DAF-2, in establishing life-stage-specific behavioral responses to CO_2_ by modulating the CO_2_-evoked activities of distinct interneurons. Finally, we identify both shared and life-stage-specific neurotransmitters and neuropeptides that regulate CO_2_ response at the two life stages. Together, our results illustrate that divergent regulatory mechanisms can establish the same chemosensory behavior through life-stage-specific changes in neural circuit function.

## Results

### Distinct sets of interneurons regulate CO_2_ response in adults vs. dauers

Well-fed *C. elegans* adults are repelled by CO_2_, whereas both starved adults and dauer larvae are attracted to CO_2_ (Hallem et al. 2011, Guillermin et al. 2017, Rengarajan et al. 2019) (Fig. S1, Fig. 1a). To elucidate the CO_2_ microcircuit of dauers, we first examined the neural activity of the CO_2_-detecting BAG neurons, which are required for CO_2_ chemotaxis in both adults and dauers (Hallem and Sternberg 2008, Hallem et al. 2011, Smith et al. 2013). We found that similar to the BAG neurons of starved adults, the BAG neurons of dauers show excitatory calcium responses to CO_2_ (Fig. S2). However, the BAG neurons of dauers showed a slightly reduced response to CO_2_ relative to the BAG neurons of starved adults, which could result from reduced diffusion of CO_2_ through the thicker cuticle of dauers or reduced sensitivity of the BAG neurons of dauers to CO_2_ (Fig. S2). Nevertheless, both starved adults and dauers show depolarizing CO_2_-evoked activity in the CO_2_-detecting sensory neurons.

**Fig 1.**
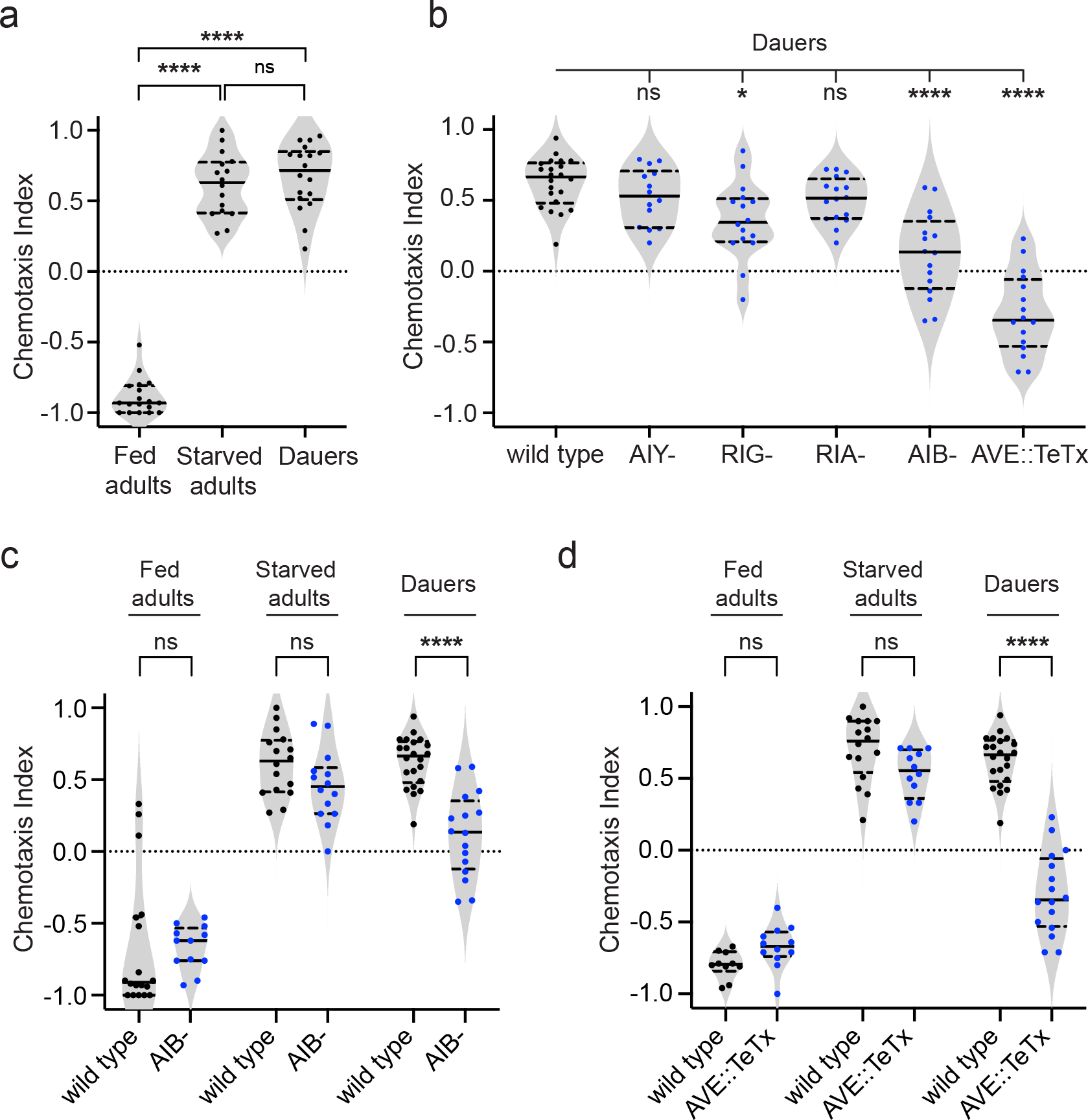
The RIG, AIB, and AVE interneurons regulate CO_2_ attraction in dauers. **a**. Well-fed adults are repelled by CO_2_, whereas starved adults and dauers are attracted to CO_2_. n = 16-18 trials per life stage and condition. \*\**\*\*p*<0.0001, ns = not significant (*p*>0.9999), Kruskal-Wallis test with Dunn’s post-test. **b.** The interneurons RIG, AIB, and AVE promote CO_2_ attraction in dauers. Strains where specific pairs of interneurons were genetically ablated (AIY-, RIG-, RIA-, or AIB-) or silenced with tetanus toxin (AVE::TeTx) were used. *\*\*\*\*p*<0.0001, *\*p*<0.05, ns = not significant (*p* = 0.8334 for wild type vs. AIY-, *p* = 0.3878 for wild type vs. RIA-), one-way ANOVA with Dunnett’s post-test. n = 14-22 trials per genotype. For a-b, each data point represents a single chemotaxis assay. Solid lines indicate medians and dotted lines indicate interquartile ranges. Responses shown are to 10% CO_2_. **c.** Behavioral responses of animals with genetically ablated AIB neurons (AIB-) in CO_2_ chemotaxis assays. \*\*\*\**p*<0.0001, ns = not significant (*p*>0.3), two-way ANOVA with Sidak’s post-test. n = 12-22 assays per life stage and condition. Each data point represents a single chemotaxis assay. Solid lines in violin plots show medians and dotted lines show interquartile ranges. Responses are to 10% CO_2_. **d.** Behavioral responses of animals expressing tetanus toxin specifically in the AVE neurons (AVE::TeTx) in CO_2_ chemotaxis assays. \*\*\*\**p*<0.0001, ns = not significant (*p*>0.06), two-way ANOVA with Sidak’s post-test. n = 10-22 assays per life stage and condition. Conventions and conditions are as in panel c.

We then investigated the CO_2_ microcircuit that operates downstream of BAG to mediate CO_2_ attraction in dauers. Although BAG neuron connectivity in dauers has not yet been characterized, in adults the BAG neurons are known to form chemical synapses with a number of interneurons, including the AIY, RIG, RIA, AIB, and AVE interneurons (Guillermin et al. 2017, Rengarajan et al. 2019). CO_2_ repulsion in well-fed adults is mediated by the AIY, RIA, and RIG interneurons (Guillermin et al. 2017). Starvation modulates the CO_2_-evoked activity of AIY and suppresses the CO_2_-evoked activity of RIG, leading to CO_2_ attraction (Rengarajan et al. 2019). To identify the interneurons required for CO_2_ attraction in dauers, we tested behavioral responses to CO_2_ in strains where individual interneurons were either genetically ablated or silenced with tetanus toxin (Guillermin et al. 2017, Katz et al. 2018). Surprisingly, we found that whereas AIY mediates CO_2_ response in both well-fed and starved adults (Guillermin et al. 2017, Rengarajan et al. 2019), it is not required for CO_2_ response in dauers: dauers lacking AIY exhibit normal CO_2_ attraction (Fig. 1b, Fig. S3). The RIA interneurons are also not required for CO_2_ attraction in dauers (Fig. 1b, Fig. S3). Instead, the RIG, AIB, and AVE interneurons mediate CO_2_ attraction in dauers. Dauers lacking RIG or AIB function showed significantly reduced CO_2_ attraction, whereas dauers lacking AVE function were repelled by instead of attracted to CO_2_ (Fig. 1b-d, Fig. S3-4). Importantly, starved adults lacking RIG function showed normal responses to CO_2_ (Rengarajan et al. 2019), and both well-fed and starved adults lacking either AIB or AVE function showed normal responses to CO_2_ (Fig. 1c-d, Fig. S4). Thus, RIG, AIB, and AVE mediate CO_2_ attraction specifically in dauers.

The only tested interneuron that was required for CO_2_ response in both dauers and adults was RIG. Although RIG is not required for CO_2_ attraction in starved adults, well-fed adults lacking RIG showed reduced CO_2_ repulsion (Guillermin et al. 2017, Rengarajan et al. 2019) and dauers lacking RIG showed reduced CO_2_ attraction (Fig. 1b, Fig. S3). Moreover, increasing neurotransmission and neuropeptide release from RIG by expressing a gain-of-function allele of the protein kinase C (*pkc-1*) gene (Sieburth et al. 2005, Sieburth et al. 2007) specifically in RIG resulted in enhanced CO_2_ repulsion in well-fed adults (Guillermin et al. 2017) and enhanced CO_2_ attraction in dauers (Fig. S5a). Thus, RIG regulates CO_2_ repulsion in well-fed adults and CO_2_ attraction in dauers. Our results demonstrate that the same interneuron can play distinct roles at different life stages.

### Similar interneuron activity promotes opposite behaviors in well-fed adults vs. dauers

To gain insight into the functional properties of the interneurons that promote CO_2_ attraction in dauers, we investigated their CO_2_-evoked calcium responses. We first examined the responses of the RIG interneurons. RIG displays excitatory CO_2_-evoked activity in well-fed adults, but this activity is silenced during starvation (Guillermin et al. 2017, Rengarajan et al. 2019). Consistent with previous findings, we observed excitatory activity in RIG in well-fed adults (Fig. 2a). Strikingly, RIG displayed similar excitatory CO_2_-evoked activity in dauers, despite the fact that well-fed adults and dauers show opposite CO_2_-evoked behaviors (Fig. 2a-b). As in adults (Guillermin et al. 2017), these excitatory responses in RIG were eliminated in dauers where the primary CO_2_-sensing BAG neurons were genetically ablated (Fig. S5b-c), indicating that RIG activity in dauers is dependent on sensory input from BAG. Together, these results suggest that the role of RIG in establishing CO_2_ response valence differs across life stages, such that excitatory activity from RIG promotes CO_2_ repulsion in well-fed adults and CO_2_ attraction in dauers.

**Fig 2.**
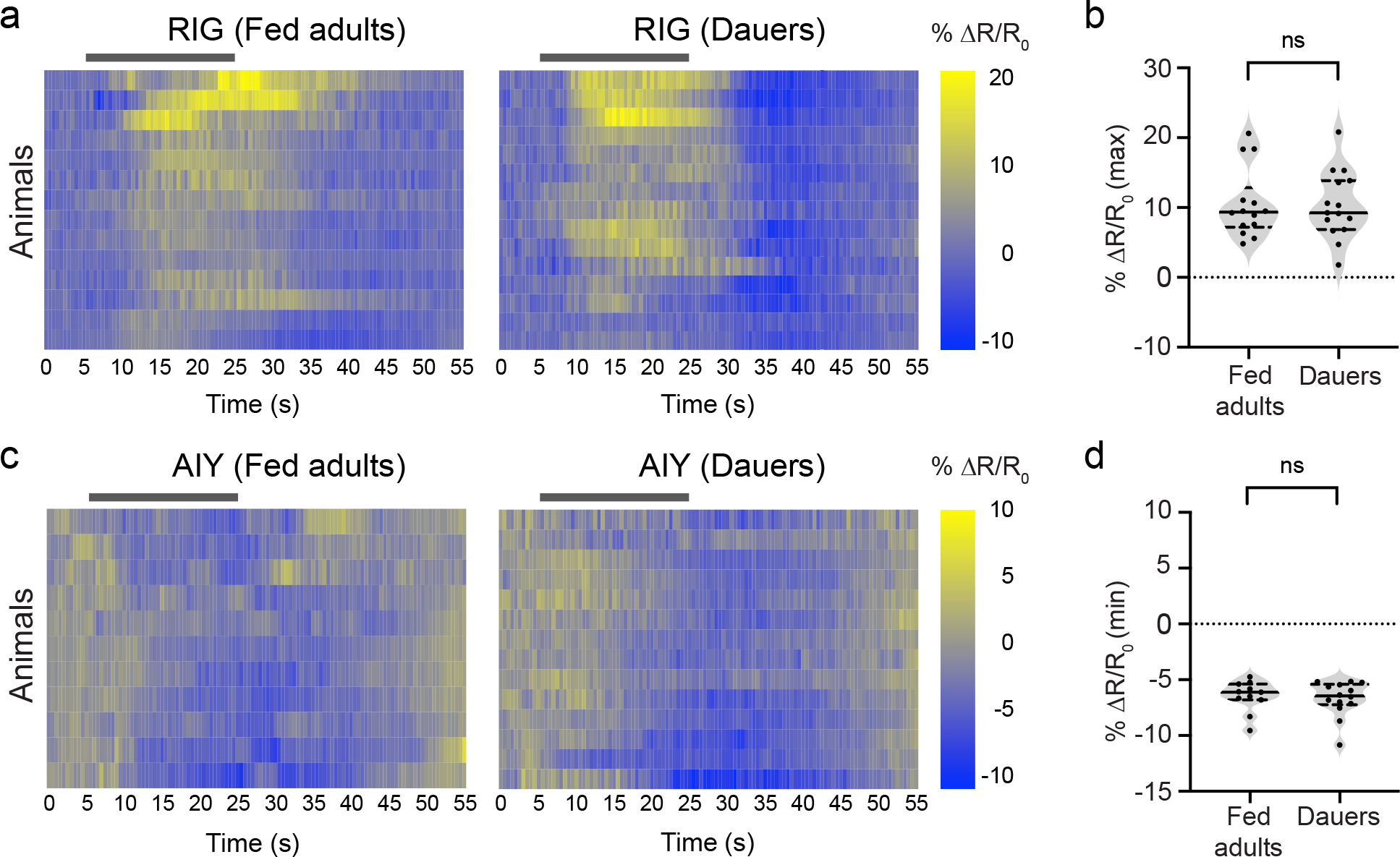
The RIG and AIY interneurons respond similarly to CO_2_ in well-fed adults and dauers. **a.** CO_2_-evoked calcium responses of RIG in well-fed adults and dauers. Each row within the heatmaps represents the response of an individual animal. Response magnitudes are color-coded according to the scale (% ΔR/R_0_) shown to the right. Responses are ordered by hierarchical cluster analysis. Gray bars indicate the timing and duration of the CO_2_ pulse. Calcium responses were measured using the ratiometric calcium indicator yellow cameleon YC3.60. Responses are to 15% CO_2_. **b.** Quantification of the maximum responses of RIG in well-fed adults and dauers. ns = not significant (*p* = 0.8989), Welch’s *t*-test. Each data point represents the response of a single animal. Solid lines in violin plots show medians and dotted lines show interquartile ranges. n = 14-15 animals per life stage. **c.** CO_2_-evoked calcium responses of AIY in well-fed adults and dauers. Conventions and conditions are as in panel a. **d.** Quantification of the minimum responses of AIY in well-fed adults and dauers. ns = not significant (*p* = 0.7377), Mann-Whitney test. Conventions are as in panel b. n = 11-14 animals per life stage.

We next examined the CO_2_-evoked calcium responses of the AIY interneurons, which are required for CO_2_ attraction in starved adults (Rengarajan et al. 2019) but not dauers (Fig. 1b, Fig. S3a). The AIY interneurons in well-fed adults are inhibited by CO_2_, and AIY ablation leads to enhanced CO_2_ avoidance (Guillermin et al. 2017). In contrast, the AIY interneurons of starved adults show stochastic responses to CO_2_ such that roughly equal proportions of animals display excitatory and inhibitory activity in AIY, and AIY-ablated starved adults show reduced CO_2_ attraction (Rengarajan et al. 2019). Thus, inhibition of AIY is associated with CO_2_ repulsion in well-fed adults and activation of AIY is associated with CO_2_ attraction in starved adults (Guillermin et al. 2017, Rengarajan et al. 2019). We found that the AIY interneurons of dauers show inhibitory calcium responses to CO_2_ that are indistinguishable from those of well-fed adults (Fig. 2c-d), despite the fact that well-fed adults and dauers exhibit opposite CO_2_-evoked behaviors. Thus, whereas starvation modulates AIY activity in adults to promote the shift from CO_2_ repulsion to CO_2_ attraction, AIY activity in dauers is the same as in well-fed adults. However, AIY appears to uncouple from the CO_2_ microcircuit in dauers such that it still responds to CO_2_ but no longer regulates CO_2_-evoked behavior. Together, these results illustrate that interneurons with similar activity patterns at different life stages may not play similar roles in regulating behavior.

### The AIB interneurons show CO_2_-evoked calcium responses specifically in dauers

In contrast to RIG and AIY, the AIB interneurons promote CO_2_ attraction selectively in dauers (Fig. 1b-c, Fig. S4). To gain a deeper understanding of how AIB regulates dauer-specific CO_2_ responses, we monitored its CO_2_-evoked calcium activity in adults and dauers. We found that AIB is mostly unresponsive to CO_2_ in well-fed and starved adults (Fig. 3a-b). In contrast, the AIB interneurons of dauers show robust excitatory calcium responses to CO_2_ (Fig. 3a-b). Thus, AIB is selectively activated by CO_2_ in dauers but not adults. AIB is known to play a role in regulating basal locomotion as well as navigation in response to multiple chemosensory cues (Gray et al. 2005, Chalasani et al. 2007, Piggott et al. 2011, Gordus et al. 2015, Summers et al. 2015). To confirm that the excitatory activity displayed by AIB in dauers was evoked by CO_2_, we measured AIB activity in response to an air control where the CO_2_ pulse was replaced with an air pulse (21% O_2_, balance N_2_) of equivalent duration. While the AIB responses of well-fed adults and starved adults were indistinguishable between CO_2_ and air controls, the CO_2_-evoked AIB responses in dauers were significantly higher than their respective air controls (Fig. S6a), suggesting that the excitatory calcium responses of AIB in dauers are evoked by CO_2_ and not due to the slight movements of animals during imaging.

**Fig 3.**
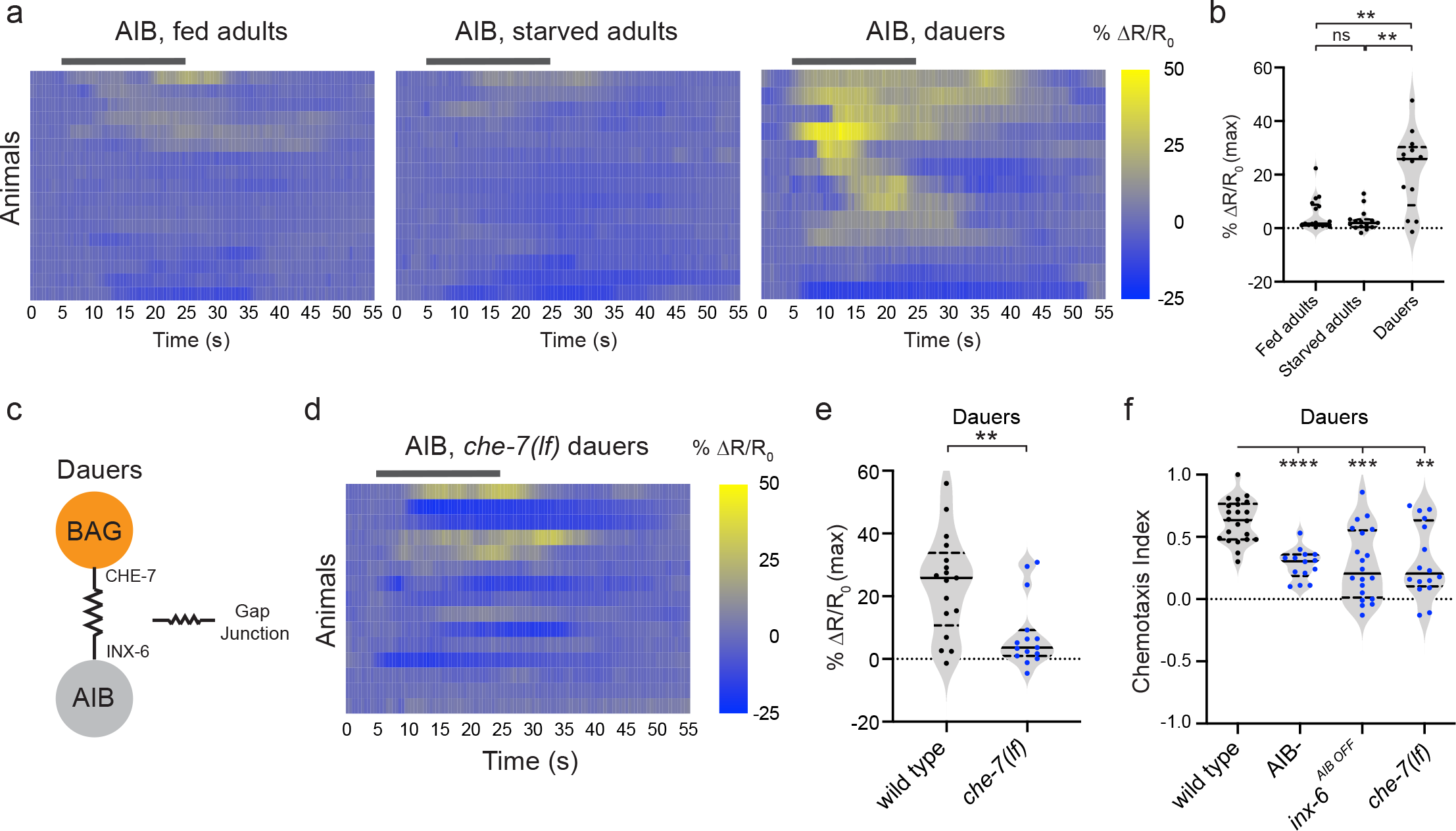
The AIB interneurons promote CO_2_ attraction specifically in dauers. **a.** The AIB interneurons show excitatory activity in dauers. Calcium responses of AIB in well-fed adults, starved adults, and dauers. Each row represents the response of an individual animal. Response magnitudes in the heatmaps are color coded according to the scale (% ΔR/R_0_) shown to the right. Responses are ordered by hierarchical cluster analysis. Gray bars indicate timing and duration of the CO_2_ pulse. Calcium responses were measured using the ratiometric calcium indicator yellow cameleon YC3.60. Responses are to 15% CO_2_. **b.** Quantification of maximum responses of AIB in well-fed adults, starved adults, and dauers. \*\**p*<0.01, ns = not significant (*p*>0.9999), Kruskal Wallis test with Dunn’s post-test. Lines in graphs show medians and interquartile ranges. n = 13-17 animals per life stage and condition. **c.** A gap junction complex is formed between the BAG sensory neurons and AIB interneurons in dauers by selective expression of the innexin subunit INX-6 in AIB. INX-6 partners with the innexin subunit CHE-7, which is expressed in BAG, to form gap junctions (Bhattacharya et al. 2019). **d.** Excitatory CO_2_-evoked calcium responses in AIB are largely eliminated in *che-7(ok2373)* mutant dauers. Conventions and imaging conditions are as in panel a. **e.** Quantification of maximum responses of AIB in wild-type and *che-7(ok2373)* dauers. Conventions and conditions are as in b. \*\**p*<0.01, Welch’s *t*-test. n = 15-17 animals per genotype. **f.** Behavioral responses of AIB-ablated (AIB-) dauers, dauers containing an AIB-specific knockout of *inx-6* (*inx-6^AIB OFF^*) (Bhattacharya et al. 2019), and *che-7(ok2373)* mutant dauers in CO_2_ chemotaxis assays. \*\**p*<0.01, \*\*\**p*<0.001, \*\*\*\**p*<0.0001, one-way ANOVA with Dunnett’s post-test. n = 14-22 trials per genotype. Each data point represents a single chemotaxis assay. Solid lines indicate medians and dotted lines indicate interquartile ranges. Responses are to 10% CO_2_.

To test whether AIB excitatory activity is required post-developmentally to establish dauer-specific CO_2_ attraction, we examined the behavioral responses of transgenic dauers with AIB-specific expression of the histamine-gated chloride channel HisCl1 (Pokala et al. 2014). We found that transient and inducible silencing of AIB by HisCl1 resulted in significantly reduced CO_2_ attraction in dauers (Fig. S6b), suggesting a non-developmental role for AIB in promoting CO_2_ attraction in dauers.

We then investigated the circuit-level changes that allow AIB to exhibit CO_2_-evoked activity specifically in dauers. The AIB interneurons form gap junctions with the BAG neurons specifically in dauers (Bhattacharya et al. 2019). Expression of the innexin gene *inx-6* specifically in AIB in dauers results in the formation of a heteromeric gap junction complex consisting of INX-6 and another gap junction subunit, CHE-7, which is expressed in BAG as well as other neurons across life stages (Bhattacharya et al. 2019) (Fig. 3c). This gap junction complex is localized at the contact points between the processes of BAG and AIB (Bhattacharya et al. 2019). To test whether the excitatory CO_2_-evoked calcium responses in AIB in dauers arise due to the dauer-specific INX-6/CHE-7 gap junction complex, we monitored CO_2_-evoked calcium activity in the AIB interneurons of *che-7* loss-of-function (*lf*) mutant dauers. We found that the strong excitatory calcium responses were almost entirely absent in *che-7(lf)* dauers (Fig. 3d-e). Moreover, in *che-7(lf)* dauers, the calcium activity of AIB in response to CO_2_ was indistinguishable from that in response to the air control (Fig. S7a). These results suggest that the CO_2_-evoked calcium responses of AIB in dauers are dependent on the dauer-specific BAG-AIB electrical synapse. In CO_2_ chemotaxis assays, *che-7(lf)* mutant dauers, as well as dauers where *inx-6(lf)* expression is eliminated specifically in AIB (*inx-6^AIB OFF^*), showed significantly reduced CO_2_ attraction compared to wild-type dauers (Fig. 3f), consistent with previous results (Bhattacharya et al. 2019). Moreover, the defects in CO_2_ attraction of *che-7(lf)* and *inx-6^AIB OFF^* dauers were comparable to those of dauers with genetically ablated AIB neurons (Fig. 3f). Ectopic expression of *inx-6* in AIB at the adult stage (Bhattacharya et al. 2019) did not affect CO_2_ responses in well-fed adults (consistent with a previous report (Bhattacharya et al. 2019)) or starved adults (Fig. S7b-c), indicating that additional mechanisms contribute to the dauer-specific role of AIB in regulating CO_2_ response. Together, these results suggest that changes in electrical synaptic wiring between the CO_2_-detecting BAG neurons and the AIB interneurons in dauers contribute to the ability of dauers to utilize a distinct CO_2_ microcircuit compared to adults.

### Inhibitory activity in a command interneuron is associated with CO_2_ attraction specifically in dauers

The AVE interneurons promote CO_2_ attraction in dauers but do not significantly alter CO_2_ responses in adults (Fig. 1b, d). To gain insight into the role of AVE in shaping CO_2_ responses specifically in dauers, we monitored CO_2_-evoked calcium responses in AVE in both adults and dauers. AVE activity is known to be associated with reversals in adults (Piggott et al. 2011), but its role in regulating chemosensory behaviors remains poorly understood. We found that AVE is mostly unresponsive to CO_2_ in well-fed adults and starved adults (Fig. 4a), consistent with our behavioral results (Fig. 1d). In contrast, AVE is predominantly inhibited by CO_2_ in dauers (Fig. 4a). The minimum peak amplitudes of the CO_2_-evoked calcium responses were significantly lower in dauers relative to adults (Fig. 4b), while the maximum peak amplitudes were similar (Fig. 4c). Thus, the AVE interneurons are specifically inhibited by CO_2_ in dauers.

**Fig 4.**
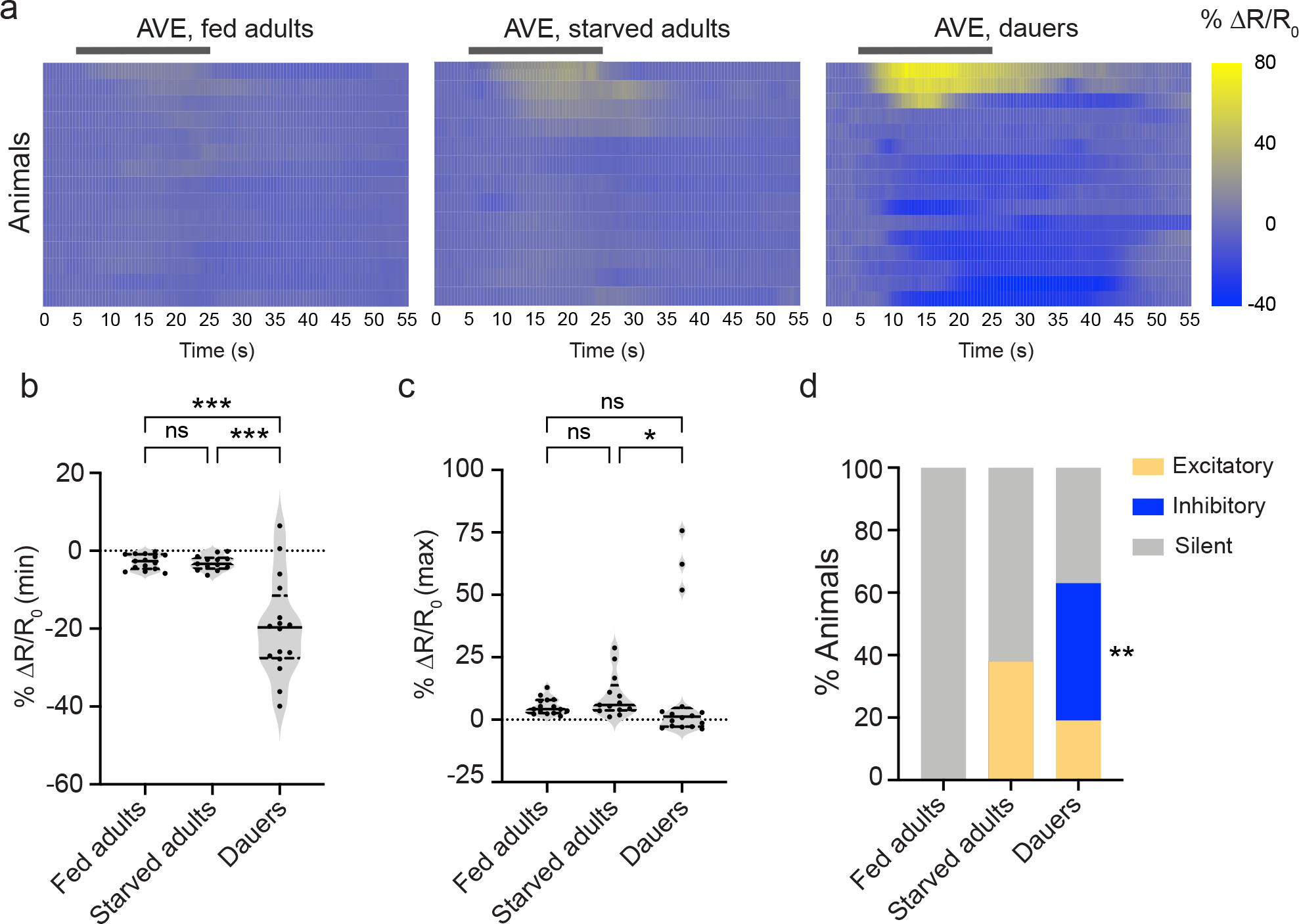
The AVE interneurons promote CO_2_ attraction specifically in dauers. **a.** Calcium responses in the AVE neurons of well-fed adults, starved adults, and dauers. Response magnitudes in the heatmaps are color-coded according to the scales (% ΔR/R_0_) shown to the right. Responses are ordered by hierarchical cluster analysis. Gray bars indicate the timing and duration of the CO_2_ pulse. Calcium responses were measured using the ratiometric calcium indicator yellow cameleon YC3.60. Responses are to 15% CO_2_. **b.** Quantification of minimum responses of AVE in well-fed adults, starved adults, and dauers. \*\*\**p*<0.001, ns = not significant (*p* = 0.9813), one-way ANOVA with Dunnett’s post-test. **c.** Quantification of maximum responses of AVE in well-fed adults, starved adults, and dauers. \**p*<0.05, ns = not significant (*p*>0.9999 for well-fed adults vs. starved adults, *p* = 0.1714 for well-fed adults vs. dauers), Kruskal Wallis test with Dunn’s post-test. In b-c, each data point represents the response of a single animal. Solid lines in violin plots show medians and dotted lines show interquartile ranges. **d.** Categorical plot displaying the percentage of excitatory, inhibitory, and silent responses in AVE across life stages and conditions. \*\**p*<0.01 for inhibitory responses of dauers vs. well-fed adults, \*\**p*<0.01 for inhibitory responses of dauers vs. starved adults, ns = not significant (*p*>0.9999) for inhibitory responses of well-fed adults vs starved adults, Fisher’s exact test. For a-d, n = 13-16 animals per life stage and condition.

To further characterize the differences in AVE activity between adults and dauers, we examined the categorical distribution of responses at the two different life stages. Responses were categorized as either excitatory or inhibitory if the absolute value of the response exceeded three standard deviations of the response to an air control (Rengarajan et al. 2019). We found that the AVE interneurons of well-fed adults were unresponsive to CO_2_ in all animals tested (Fig. 4d). Starved adults showed excitatory AVE responses in ∼39% of animals tested; the AVE neurons of the remaining animals were unresponsive to CO_2_ (Fig. 4d). However, the AVE responses we observed in starved adults exposed to CO_2_ were not significantly different from the AVE responses of starved adults exposed to an air control, indicating that these responses were not specifically evoked by CO_2_ (Fig. S8a-b). In contrast, dauers showed inhibitory responses in AVE in roughly 45% of animals, a pattern that we did not observe in adults (Fig. 4d). In addition, the inhibitory AVE responses we observed in dauers were significantly different between CO_2_ and air controls (Fig. S8b), indicating that these responses were evoked by CO_2_. Although a smaller percentage (20%) of dauers showed excitatory activity in AVE, the excitatory responses of AVE to CO_2_ were not significantly different from those observed in response to the air control (Fig. S8a), indicating that they were not CO_2_-evoked. Moreover, the excitatory AVE responses of dauers did not differ significantly from those of well-fed or starved adults (Fig. 4d). Together, our results indicate that AVE command interneurons show CO_2_-evoked inhibitory activity specifically in dauers.

### Insulin signaling establishes life-stage-specific CO_2_ responses by modulating the activities of distinct interneurons in dauers vs. adults

We next investigated the molecular mechanisms that shape CO_2_ response across life stages. We first examined the role of insulin signaling in regulating CO_2_ response in dauers and adults, since the insulin pathway plays an important role in regulating the developmental decision to enter the dauer state (Hu 2007). Insulin signaling also regulates a number of chemosensory behaviors in adults, including salt chemotaxis (Tomioka et al. 2006, Adachi et al. 2010) and acute CO_2_ avoidance (Hallem and Sternberg 2008). However, its role in regulating chemosensory responses in dauers had not been examined. We first examined the CO_2_ response of animals carrying a loss-of-function mutation in the *daf-2* gene, which encodes the sole *C. elegans* homolog of the mammalian insulin/IGF receptor (Hu 2007). We found that both well-fed and starved *daf-2(lf)* mutant adults were strongly attracted to CO_2_ (Fig. 5a), reflecting a switch in CO_2_ response valence in well-fed adults. In contrast, *daf-2(lf)* mutant dauers were unresponsive to CO_2_ (Fig. 5a). In addition, we found that dauers carrying loss-of-function mutations in the 3-phosphoinositide-dependent-kinase-1 gene *pdk-1* and the protein kinase B (Akt/PKB) gene *akt-1*, both of which act downstream of *daf-2* in the *C. elegans* insulin pathway (Hu 2007), showed significantly reduced CO_2_ attraction (Fig. S9a). Thus, insulin signaling plays a life-stage-specific role in modulating CO_2_ response: it promotes CO_2_ repulsion in well-fed adults and CO_2_ attraction in dauers.

**Fig 5.**
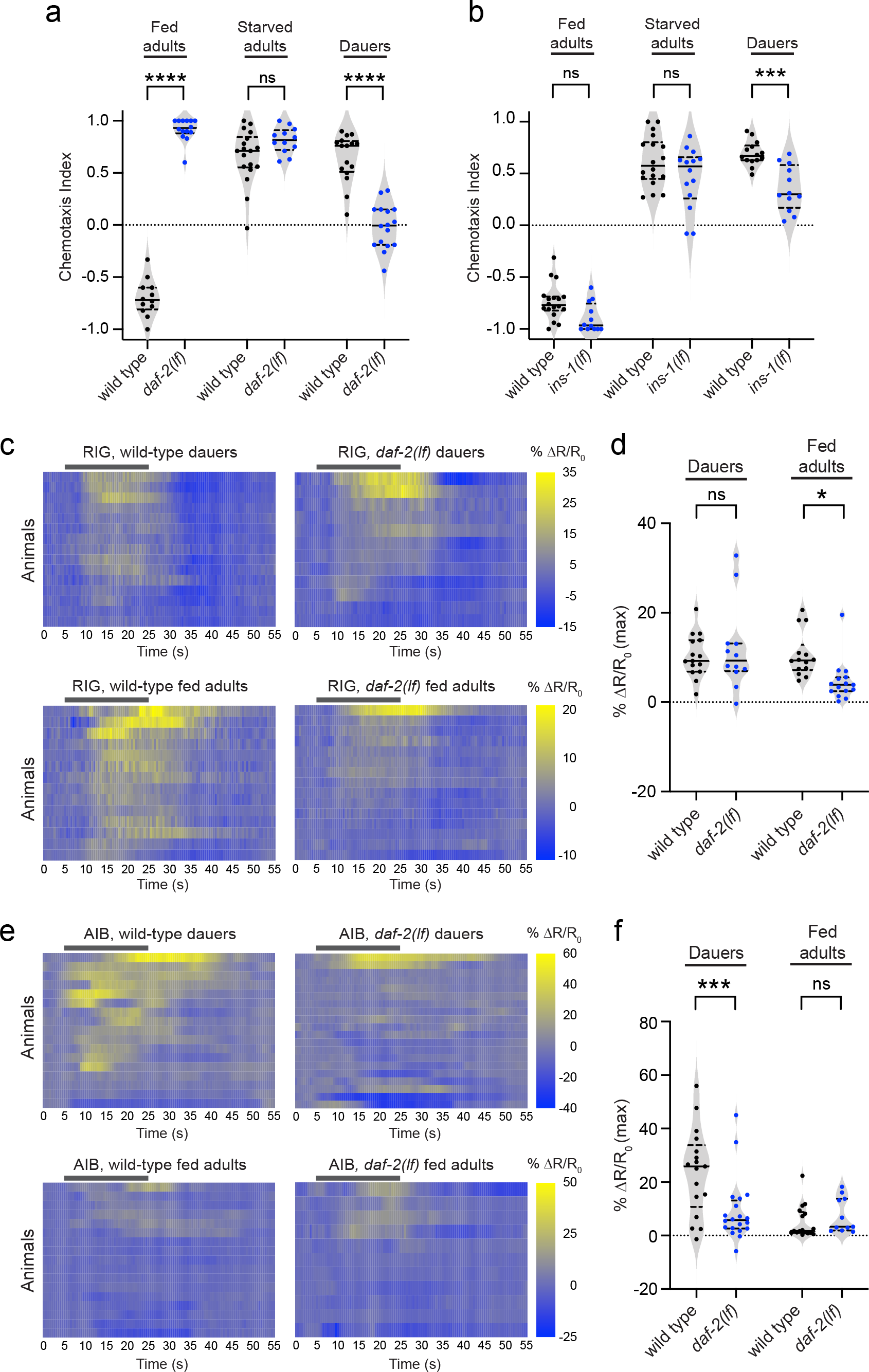
Insulin signaling establishes CO_2_ responses across life stages. **a.** DAF-2 establishes CO_2_ repulsion in well-fed adults and CO_2_ attraction in dauers. \*\*\*\**p*<0.0001, ns = not significant (*p* = 0.1202), two-way ANOVA with Sidak’s post-test. n = 12-18 assays per life stage and condition. **b.** INS-1 promotes CO_2_ attraction in dauers. \*\*\**p*<0.001, ns = not significant (*p*>0.1), two-way ANOVA with Sidak’s post-test. n = 12-18 assays per life stage and condition. For a-b, graphs show the behavioral responses of *daf-2(e1370)* and *ins-1(nr2091)* mutants across life stages and conditions in CO_2_ chemotaxis assays. Each data point represents a single chemotaxis assay. Solid lines in violin plots show medians and dotted lines show interquartile ranges. Responses are to 10% CO_2_. **c.** CO_2_-evoked calcium responses of the RIG neurons of wild-type and *daf-2(e1370)* mutant well-fed adults and dauers. Response magnitudes in the heatmaps are color-coded according to the scales (% ΔR/R_0_) shown to the right. Responses are ordered by hierarchical cluster analysis. Gray bars indicate the timing and duration of the CO_2_ pulse. Calcium responses were measured using the ratiometric calcium indicator yellow cameleon YC3.60. Responses are to 15% CO_2_. **d.** Quantification of the maximum responses of RIG in wild-type and *daf-2(e1370)* mutant dauers and well-fed adults. \**p*<0.05, ns = not significant (*p* = 0.7649), two-way ANOVA with Sidak’s post-test. n = 12-15 animals per life stage and genotype. **e.** CO_2_-evoked calcium responses of the AIB neurons of wild-type and *daf-2(e1370)* mutant dauers and well-fed adults. Conventions and conditions are as in panel c. **f.** Quantification of the maximum responses of AIB in wild-type and *daf-2(e1370)* mutant well-fed adults and dauers. Conventions are as in panel d. \*\*\**p*<0.001, ns = not significant (*p* = 0.8081), two-way ANOVA with Sidak’s post-test. n = 11-20 animals per life stage and genotype.

We next sought to identify the DAF-2 ligands that modulate CO_2_ response. The *C. elegans* genome encodes 40 insulin-like peptides (ILPs) (Li and Kim 2008). To identify ILPs that may be involved in establishing CO_2_ attraction in dauers, we examined a subset of ILP genes known to be transcriptionally upregulated upon dauer entry (Lee et al. 2017). We found that dauers carrying loss-of-function mutations in the *ins-1* gene showed significantly reduced CO_2_ attraction compared to wild-type dauers (Fig. 5b, Fig. S9b). In contrast, CO_2_ repulsion in well-fed adults and CO_2_ attraction in starved adults were unaffected in *ins-1(lf)* mutants (Fig. 5b). These results strongly suggest that the modulatory actions of INS-1 are specific to dauers, although we cannot exclude the possibility that INS-1 may act redundantly with other peptides to regulate CO_2_ response in adults.

To better understand how insulin signaling establishes life-stage-specific CO_2_ responses, we examined the effects of DAF-2 on neural activity within the CO_2_ microcircuit. We first examined CO_2_-evoked calcium responses of RIG in *daf-2(lf)* mutants, since RIG excitatory activity was associated with opposite behavioral responses to CO_2_ in well-fed adults vs. dauers (Fig. 2a-b). We found that RIG showed similar excitatory calcium responses in *daf-2(lf)* mutant dauers compared to wild-type dauers (Fig. 5c-d). In contrast, RIG was largely silent in *daf-2(lf)* mutant well-fed adults (Fig. 5c-d). These results suggest that DAF-2 modulates RIG activity specifically in well-fed adults. We previously showed that activation of RIG promotes CO_2_ repulsion in well-fed adults, and silencing RIG accelerates the shift from CO_2_ repulsion to CO_2_ attraction during starvation (Guillermin et al. 2017, Rengarajan et al. 2019). Thus, the CO_2_ attraction observed in *daf-2(lf)* well-fed adults likely arises in part from the loss of CO_2_-evoked activity in RIG.

We next examined CO_2_-evoked calcium responses in AIB, since AIB promotes CO_2_ attraction specifically in dauers (Fig. S4). We found that the excitatory CO_2_-evoked calcium responses in AIB seen in wild-type dauers were largely eliminated in *daf-2(lf)* dauers (Fig. 5e-f). These results suggest that the dauer-specific calcium responses of AIB are dependent on insulin signaling. In contrast, the AIB responses of *daf-2(lf)* well-fed adults were indistinguishable from those of wild-type adults; neither displayed CO_2_-evoked activity (Fig. 5e-f). Thus, the loss of CO_2_ attraction in *daf-2(lf)* dauers likely arises in part from the silencing of AIB activity. Together, our results suggest that insulin signaling establishes life-stage-specific CO_2_ responses by fine-tuning the activities of distinct interneurons within the CO_2_ microcircuit.

### Both shared and distinct molecular signals establish CO_2_ response in adults vs. dauers

We next investigated additional signaling mechanisms that may establish CO_2_ response in adults vs. dauers. The CO_2_-detecting BAG neurons express the vesicular glutamate transporter EAT-4 (Serrano-Saiz et al. 2013), and well-fed adults carrying a loss-of-function mutation in the *eat-4* gene show dramatically reduced CO_2_ repulsion (Guillermin et al. 2017). We found that *eat- 4(lf)* dauers showed normal responses to CO_2_, while starved *eat-4(lf)* adults showed significantly reduced CO_2_ attraction (Fig. 6a). Thus, glutamate signaling is required for CO_2_ response in adults but not dauers. We next investigated the CO_2_ response of dauers carrying a loss-of-function mutation in the tyrosine decarboxylase gene *tdc-1*, which regulates tyraminergic and octopaminergic signaling (Chase and Koelle 2007), since TDC-1 promotes CO_2_ attraction during starvation in adults (Rengarajan et al. 2019). We found that, similar to CO_2_ attraction in starved adults, CO_2_ attraction in dauers was significantly reduced in *tdc-1(lf)* mutants (Fig. 6b). These results suggest that both dauers and starved adults utilize biogenic amine signaling to promote CO_2_ attraction.

**Fig 6.**
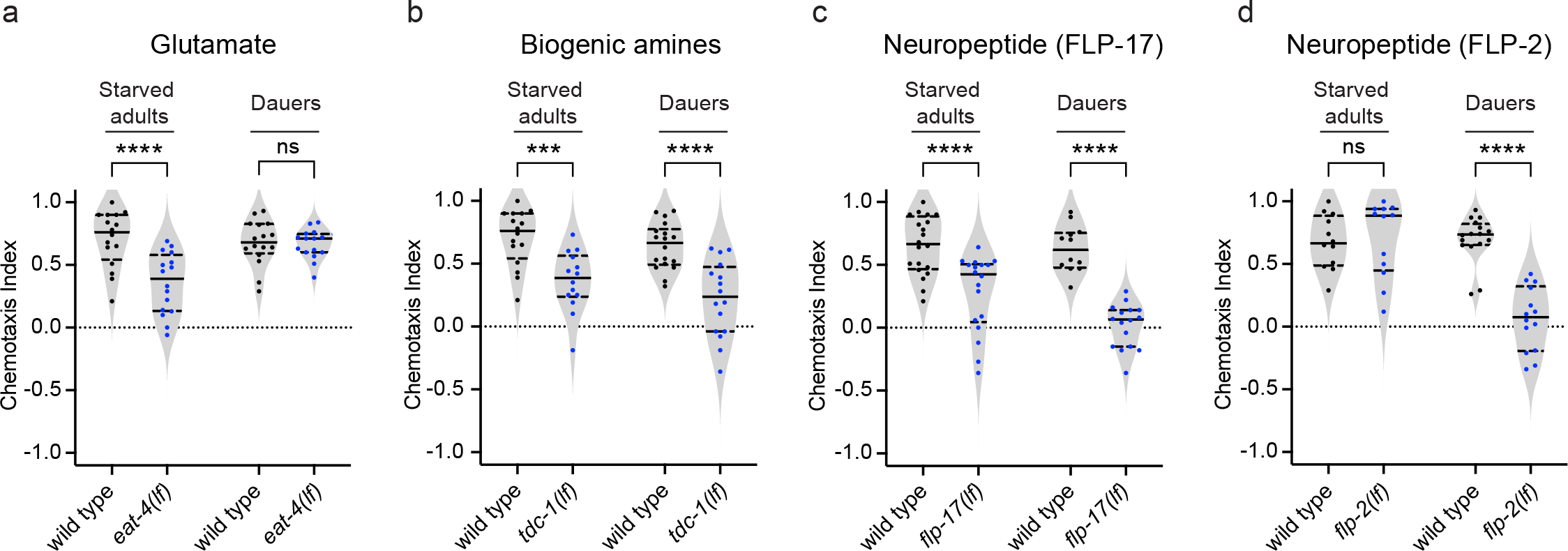
Both shared and distinct molecular signals establish CO_2_ attraction in dauers and starved adults. Behavioral responses of starved adults and dauers containing loss-of-function mutations in the vesicular glutamate transporter gene *eat-4* (**a**), the tyrosine decarboxylase gene *tdc-1* (**b**), the neuropeptide gene *flp-17* (**c**), or the neuropeptide gene *flp-2* (**d**) in CO_2_ chemotaxis assays. \*\*\*\**p*<0.0001, \*\*\**p*<0.001, ns = not significant (*p*>0.9), two-way ANOVA with Sidak’s post-test. n = 12-20 assays per life stage and genotype. Each data point represents a single chemotaxis assay. Solid lines in violin plots show medians and dotted lines show interquartile ranges. Responses are to 10% CO_2_.

Neuropeptide signaling plays a critical role in modulating neural circuit function and behavior across species (Li and Kim 2008), and was previously shown to modulate the CO_2_-evoked behavioral responses of *C. elegans* adults (Guillermin et al. 2017). To examine whether CO_2_ responses in dauers are regulated by neuropeptides, we performed a candidate screen of dauers carrying loss-of-function mutations in 16 neuropeptide genes (Fig. S10a). We focused on neuropeptide genes that are either expressed in BAG neurons (Hallem et al. 2011); transcriptionally upregulated upon dauer entry (Lee et al. 2017); known to play a role in regulating CO_2_ response in adults (Guillermin et al. 2017, Rengarajan et al. 2019); or that encode neuropeptides dependent on the neuropeptide-processing gene *sbt-1,* since *sbt-1* promotes dauer entry and regulates CO_2_ attraction in dauers (Lee et al. 2017). We found that dauers carrying loss-of-function mutations in the BAG-expressed FMRFamide-like neuropeptide gene *flp-17* showed significantly reduced CO_2_ attraction compared to wild-type dauers (Fig. S10a, Fig. 6c). FLP-17 was previously shown to be required for CO_2_ repulsion in well-fed adults (Guillermin et al. 2017); we found that it is also required for normal CO_2_ attraction in starved adults (Fig. 6c). Thus, FLP-17 promotes CO_2_ response in both dauers and adults, regardless of CO_2_ response valence.

We also found that dauers with loss-of-function mutations in the FMRFamide-like neuropeptide gene *flp-2* showed significantly reduced CO_2_ attraction (Fig. S10, Fig. 6d), while CO_2_ repulsion in well-fed adults and CO_2_ attraction in starved adults remained unaffected in *flp-2* mutants (Fig. S10b, Fig. 6d). Thus, FLP-2 neuropeptides play a novel role in regulating CO_2_ attraction specifically in dauers. In contrast, we found that *nlp-1* and *flp-16*, which were previously shown to regulate CO_2_ attraction in starved adults (Rengarajan et al. 2019), did not regulate CO_2_ attraction in dauers (Fig. S10a). Thus, NLP-1 and FLP-16 play adult-specific roles in regulating CO_2_ response. Together, our results demonstrate that both shared and life-stage-specific molecular pathways drive CO_2_-evoked behaviors across life stages. Moreover, the similar behavioral responses of dauers and starved adults are shaped by both shared and distinct molecular mechanisms.

## Discussion

We have shown that dauers and starved adults achieve the same behavioral state, CO_2_ attraction, using distinct neural circuits. While the same CO_2_-detecting sensory neurons are required for CO_2_ response in dauers and adults, distinct sets of interneurons mediate these responses (Fig. 7). Some interneurons are required for CO_2_ response in adults but not dauers, some are required in dauers but not adults, and some promote opposite responses to CO_2_ at the two life stages (Fig. 7). Thus, the same behavior can arise from distinct patterns of interneuron activity.

**Fig 7.**
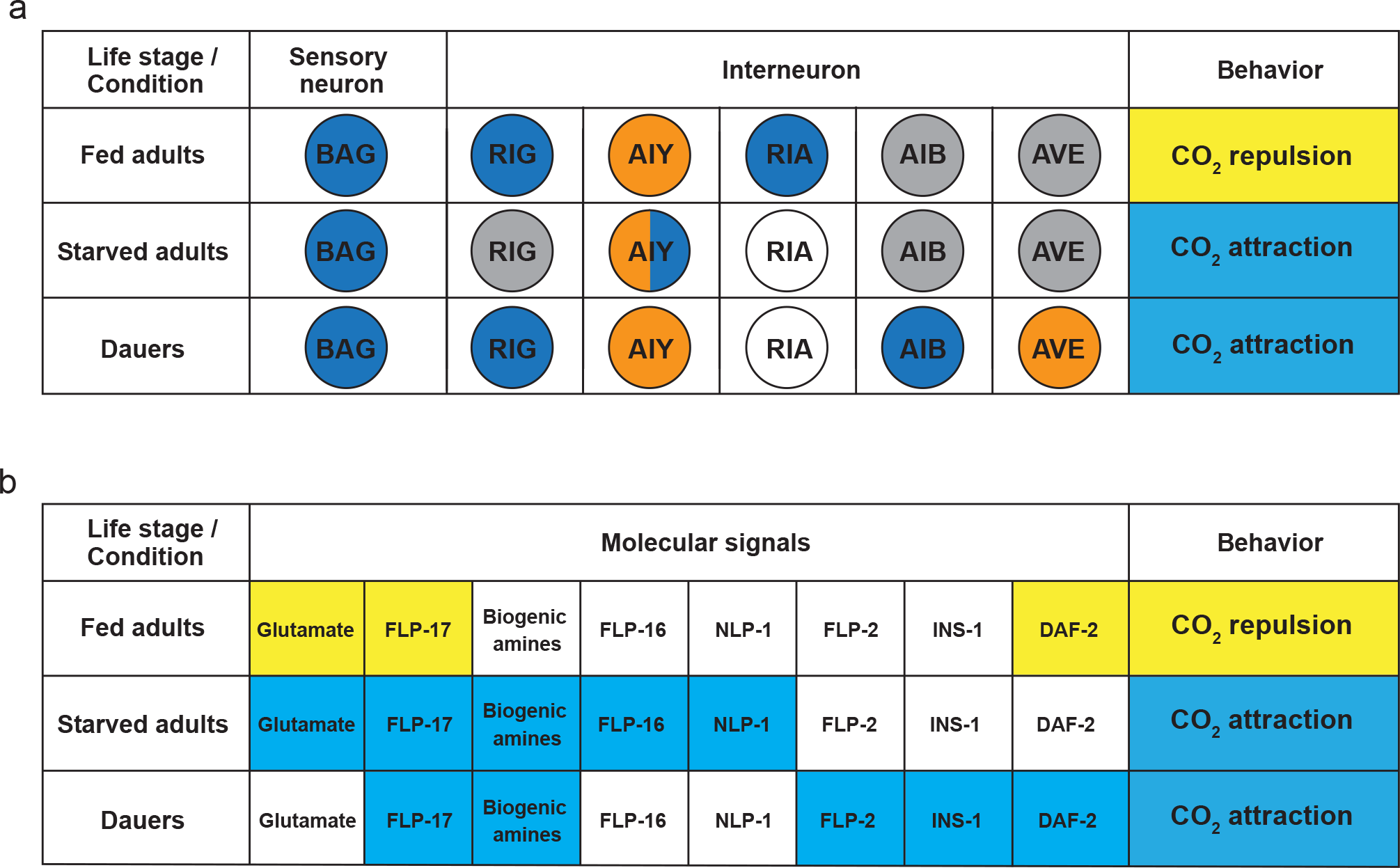
Distinct mechanisms shape CO_2_ response across life stages. **a.** Table showing the CO_2_-evoked activity of the BAG sensory neurons and downstream interneurons associated with CO_2_ repulsion (yellow) in well-fed adults and CO_2_ attraction (light blue) in starved adults and dauers. Excitatory and inhibitory neural activity is indicated by dark blue and orange shading, respectively. Non-responsive neurons are indicated by gray shading; neurons whose CO_2_-evoked calcium activity has not been examined because the neurons are not required for CO_2_-evoked behavior are indicated by white shading. For AVE interneuron, color codes indicate predominant calcium responses across life stages. **b.** Table showing the requirement for different molecular signals in establishing CO_2_ response across life stages. Neurotransmitters and neuropeptides required to establish CO_2_ repulsion and CO_2_ attraction are indicated by yellow and blue shading, respectively. The lack of requirement for a particular molecular signal in establishing CO_2_ response is indicated by white shading.

Previous studies in *C. elegans* have shown that the same thermotaxis behavior can result from the activities of different combinations of thermosensory neurons in different environmental contexts as a result of sensory neuron degeneracy (Beverly et al. 2011, Yeon et al. 2021). In contrast to this scenario, the same pair of sensory neurons is required for CO_2_ response across contexts and life stages (Hallem et al. 2011, Guillermin et al. 2017, Rengarajan et al. 2019). Differences in the CO_2_ circuit composition of dauers vs. adults arise instead at the level of interneurons and do not reflect circuit degeneracy (Fig. 7). Although it is unclear why dauers utilize a distinct set of interneurons from adults to mediate CO_2_ response, one possibility is that the dauer circuit reflects the need for dauers to coordinate CO_2_ attraction with other dauer-specific behaviors, such as nictation and dauer recovery (Hu 2007). It remains to be determined if CO_2_ acts as a sensory cue to drive these or other dauer-specific behaviors.

While most interneurons in the CO_2_ microcircuit are required for CO_2_ response exclusively in adults or dauers, the RIG interneurons are required for CO_2_ response at both life stages (Guillermin et al. 2017, Rengarajan et al. 2019) (Fig. 1b). Surprisingly, however, excitatory CO_2_-evoked activity in RIG is associated with opposite behavioral responses in well-fed adults vs. dauers: the depolarizing activity of RIG promotes CO_2_ repulsion in adults and CO_2_ attraction in dauers (Fig. 2a-b). Thus, the “meaning” of RIG activity varies across life stages. While the connectomes of adults and non-dauer larvae are known (White et al. 1986, Witvliet et al. 2020), the connectome of dauer larvae has not yet been mapped. Thus, whether the meaning of RIG activity changes as a result of dauer-specific structural changes in neural connectivity or functional changes in signaling remains to be determined.

The AIB interneurons contribute to CO_2_ attraction specifically in dauers at least in part as a result of dauer-specific gap junctions between BAG and AIB (Bhattacharya et al. 2019) (Fig. 3). CO_2_-evoked activity occurs in AIB specifically in dauers and requires the CHE-7/INX-6 gap junction complex (Fig. 3, Fig. S6b). The BAG neurons form a chemical synapse with AIB in adults (White et al. 1986, Witvlet et al. 2020), but the presence of this synapse does not appear to be sufficient for CO_2_-evoked activity in the AIB neurons (Fig. 3a-b). It is possible that the electrical synapses between BAG and AIB in dauers lead to alterations in the composition and/or function of the chemical synapses between BAG and AIB, as has been shown for a different synapse in *C. elegans* (Liu et al. 2017). We have also shown that dauer-specific AIB activity is dependent on insulin signaling (Fig. 5e-f). Thus, AIB activity in dauers is shaped by the combined actions of the insulin pathway and gap junction signaling.

Together, our results demonstrate that CO_2_ attraction in dauers vs. adults is established via distinct neural circuits and regulated by shared as well as distinct molecular pathways (Fig. 7). In future studies, it will be interesting to determine whether different neural and molecular mechanisms also underlie similar chemosensory behaviors in other organisms, including humans. In addition, dauer larvae are developmentally similar to the infective larvae of parasitic nematodes (Crook 2014), which infect over one billion people worldwide and cause some of the most devastating neglected tropical diseases (Schafer and Skopic 2006, Lustigman et al. 2012). The infective larvae of multiple parasitic nematode species use CO_2_ as a host-seeking cue (Hallem et al. 2011, Castelletto et al. 2014, Lee et al. 2016, Ruiz et al. 2017, Banerjee and Hallem 2020), but the neural mechanisms that drive these responses remain unknown. Thus, a better understanding of how *C. elegans* responds to CO_2_ may lead to new strategies for controlling these devastating parasites.

## Methods

### *C. elegans* strains

Worms were cultured and maintained on 2% Nematode Growth Media (NGM) plates seeded with *Escherichia coli* OP50 bacteria at ambient temperature (∼22°C) and CO_2_ (∼0.038%) following standard procedures (Stiernagle 2006, Scott 2011). The temperature-sensitive, dauer-constitutive strains CB1370, EAH382, EAH383 and JT191 were maintained at 15°C but were moved to ambient temperature (∼22°C) at least 24 h prior to experiments to minimize any effects of temperature shifts on behavior or neural calcium responses. Notably, temperature shifts performed on the wild-type (N2) strain did not significantly alter behavioral responses in CO_2_ chemotaxis assays (*p*>0.6, Welch’s t-test). Strains used in this study are listed in Extended Data Table 1. The strains EAH381 and EAH383 were generated by crossing EAH259 *Ex[odr-2(1b)::YC3.60; lin-44::GFP]* to RB1834 *che-7(ok2373)* and CB1370 *daf-2(e1370)*, respectively. The strain EAH382 was generated by crossing PS6028 *syEx1134[twk-3::YC3.60; pax-2::GFP]* to CB1370 *daf-2(e1370)*.

### Preparation of animals for CO_2_ chemotaxis assays

#### Well-fed adults

All experiments were performed with young adults (∼1 day old) as previously described (Guillermin et al. 2017, Rengarajan et al. 2019). Briefly, animals were collected in a 65 mm Syracuse watch glass by washing them off plates with M9 buffer. Animals were washed twice with M9 buffer and then once with ddH_2_O in the watch glass. Animals were then gently transferred onto a small rectangular piece of Whatman filter paper, which was used to transport them to the center of a 10 cm 2% NGM plate without food for chemotaxis assays.

#### Starved adults

Young adults were washed in a watch glass as described above, and then starved on a 10 cm 2% NGM plate without bacterial food for 3 h as previously described (Rengarajan et al. 2019). Animals were placed within an annular ring of filter paper soaked in 20 mM copper chloride (CuCl_2_) solution to keep them from crawling off the edges of plates, since copper is aversive to *C. elegans* (Campbell et al. 2017). After 3 h of starvation, the CuCl_2_ ring was removed, and the animals were collected from the plate and washed twice in M9 and once in ddH_2_O in a watch glass. Animals were then transferred onto the center of a 10 cm 2% NGM plate for chemotaxis assays using a piece of Whatman filter paper.

#### Dauers

To generate dauer larvae, 8-10 young adults were transferred to 2% NGM plates with a lawn of OP50 and left for 10-14 days at room temperature until the OP50 on the plates was depleted. Dauers were differentiated from other life stages on the basis of resistance to sodium dodecyl sulphate (SDS), as described previously (Karp 2016). Briefly, animals were washed off food-depleted plates with dH_2_O into a 15 mL conical tube and centrifuged at 1000 rpm for 60-90s. The supernatant was discarded without disturbing the worm pellet. The tube was then filled with 5 mL 1% SDS solution and gently mixed on a rotator at room temperature for 15 min. Following SDS treatment, the tube was centrifuged at 1000 rpm for 60-90 s, and the SDS was removed without disturbing the pellet. 10 mL dH_2_O was then added to the tube, which was mixed thoroughly and then centrifuged at 1000 rpm for 60-90 s. Washes with 10 mL dH_2_O were repeated twice for a total of three washes. Finally, dauers were transferred in water drops onto 2% NGM plates for chemotaxis assays as indicated (Fig. S1).

### CO_2_ chemotaxis assays

CO_2_ chemotaxis assays were performed as previously described (Guillermin et al. 2017, Rengarajan et al. 2019). Animals were placed onto the center of a 10 cm 2% NGM plate at the start of assay. The CO_2_ stimulus (the desired concentration of CO_2_, 21% O_2_, balance N_2_) and air stimulus (21% O_2_, balance N_2_) were pumped through holes in opposite sides of the plate lid to establish a CO_2_ gradient (Fig. S1a). Gas stimuli were delivered using a syringe pump (PHD 2000, Harvard Apparatus) through ¼-inch flexible PVC tubing at flow rates of 2 mL/min (for adult assays) or 0.5 mL/min (for dauer assays). Assays ran for 20 min (for adults) or 1 h (for dauers). At the end of each assay, the number of animals within a 20 mm diameter circle under each gas inlet (in the case of adults) or within the indicated 30 mm segments on both sides of the plate (in the case of dauers) were counted (Fig. S1b-c). For transgenic strains containing extrachromosomal arrays, only the animals expressing fluorescent markers within the scoring regions were counted under a fluorescent dissecting microscope. A Chemotaxis Index (CI) was then calculated as:

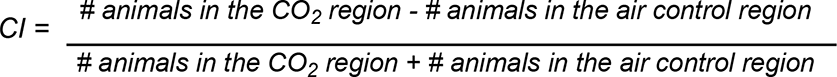

To control for directional bias due to room vibration or other sources, assays were conducted in pairs, with the CO_2_ gradients in opposite orientations for the two plates. If the absolute value of the difference in CI between two assays in a pair was ≥0.9, both assays were discarded as behavior was assumed to be impacted by directional bias. Assays were also discarded if fewer than 7 animals moved to the combined scoring regions. For strain RB2575, which showed decreased motility, single assays within a pair that had more than 7 animals in the combined regions were scored and included in the analysis even if fewer than 7 animals moved in the other assay in the pair, provided there was no directional bias within the pair. In the case of the neuropeptide genes *ins-1*, *flp-2,* and *flp-17*, two independent alleles of each gene were tested in CO_2_ chemotaxis assays. The alleles *nr2091* and *nj32* of the *ins-1* gene were used in Fig. 5b and Fig. S9b, respectively. The alleles *gk1039* and *ok3351* of the *flp-2* gene were used in Fig. 6d and Fig. S10a, respectively. The alleles *n4894* and *ok3587* of the *flp-17* gene were used in Fig. 6c and Fig. S10a, respectively.

### Histamine assays with dauers

Transgenic dauers expressing the histamine-gated chloride channel HisCl1 (Pokala et al. 2014) were isolated using 1% SDS treatment as described above. Dauers were then incubated in 20 mM histamine solution (in dH_2_O) for 1 h prior to assays. For no-histamine control experiments, dauers were incubated in dH_2_O without histamine for the same duration. Dauers were then transferred onto 2% NGM plates without bacteria, with or without 20 mM histamine, and CO_2_ chemotaxis assays were performed as described above.

### Calcium imaging

Calcium imaging experiments were performed as previously described (Carrillo et al. 2013, Guillermin et al. 2017, Rengarajan et al. 2019) using the genetically encoded calcium indicator yellow cameleon YC3.60 (Nagai et al. 2004). To starve animals prior to imaging, young adults expressing YC3.60 were picked from NGM plates with OP50 to NGM plates without bacteria and allowed to crawl for 1 min to remove residual bacteria. Animals were then transferred to 2% NGM plates (without food) lined with an annular ring of CuCl_2_-soaked Whatman paper and starved for 3-6 h before imaging. Dauers for imaging were isolated by SDS treatment as described above.

For imaging, adults were immobilized onto a freshly made 2% agarose pad (made with 10 mM HEPES) on a cover glass using Meridian Surgi-Lock 2oc surgical glue. Dauers were immobilized onto a dry 2% agarose pad (in 10 mM HEPES) on a cover glass since dauers are resistant to glue. A chamber fabricated from a 6 cm Petri dish with a 15 mm hole at the base and two 5 mm holes on diametrically opposite sides for gas inlets was placed around the animal and secured onto the cover glass with beeswax. The chamber was humidified using wet tissue wipes. Gases were delivered into the chamber at flow rates of 0.7-0.8 L/min (controlled by a flow meter) from two gas tanks fitted with valves controlled by a ValveBank TTL pulse generator. An air pulse (21% O_2_, balance N_2_) was delivered for 20 s, followed by a 20 s CO_2_ pulse (15% CO_2_, 21% O_2_, balance N_2_) and then another air pulse (21% O_2_, balance N_2_) for 30 s. For air controls, the CO_2_ pulse was replaced with an air pulse of equivalent duration (20 s). Imaging was performed using a Zeiss AxioObserver A1 inverted microscope equipped with a 40x EC Plan-NEOFLUAR lens and a Hamamatsu C9100 EM-CCD camera. Images were acquired in the YFP and CFP channels at 2 frames/s using Zeiss AxioVision software.

Image analyses were performed using AxioVision software and Microsoft Excel. Images were analyzed by selecting regions of interest (ROIs) that consisted of either the soma (BAG, RIG, AVE) or process (AIY, AIB) of the neuron of interest and a background region. For AIY imaging, the synapse-rich part of the process previously designated as zone 2 (Colon-Ramos et al. 2007) was selected as the ROI. For AIB imaging, a segment of the process where spatiotemporal expression of INX-6 was observed in dauers (Bhattacharya et al. 2019) was selected as the ROI. The average intensity for YFP and CFP of the background region was subtracted from the average intensity for YFP and CFP of the soma or process. YFP values were adjusted to correct for CFP signal bleed-through, and the YFP/CFP ratio was then calculated. The data were baseline-adjusted for linearity using the 10 s before and after the gas stimulus as the baseline. For each dataset, the different genotypes, life stages, or conditions were tested in parallel.

For quantification, the response period was defined as the time interval beginning with the onset of the CO_2_ pulse and ending 10 s after the offset of the CO_2_ pulse. The % ΔR/R_0_ (max) and % ΔR/R_0_ (min) values were calculated during the response period. For AVE imaging, categorization of responses as excitatory, inhibitory, or silent was performed using threshold values generated from air control experiments performed for each life stage or condition. Maximum and minimum threshold values were set as 3 standard deviations above the mean % ΔR/R_0_ (max) air response or 3 standard deviations below the mean % ΔR/R_0_ (min) air response, respectively. A CO_2_ response with a % ΔR/R_0_ (max) value higher than the maximum threshold value was designated as excitatory; a response with a % ΔR/R_0_ (min) value lower than the minimum threshold value was designated as inhibitory. A CO_2_ response where the most extreme % ΔR/R_0_ values were within the maximum and minimum thresholds was designated as silent. For all recordings except RIG, recordings were excluded from quantification if one or more of the % ΔR/R_0_ values during the 5 s interval preceding CO_2_ onset was outside of the range defined by ±3 standard deviations from the mean % ΔR/R_0_ (max) or % ΔR/R_0_ (min) values of an air stimulus during the same time interval (air stimuli were delivered to separate sets of worms in control experiments). Heatmaps were generated using GraphPad Prism v9.1.0. Responses within heatmaps were ordered by hierarchical clustering analysis using the web-based tool Heatmapper (Babicki et al. 2016), using Euclidean distance as a similarity measure.

### Statistical analysis

Statistical tests were performed using GraphPad Prism v9.1.0. Specific statistical tests used are indicated in the figure legends. Normality was determined using a D’Agostino-Pearson omnibus normality test; if data were not normally distributed, non-parametric tests were used. Power analyses were performed using G*Power v3.1.9.6 (Faul et al. 2007). We note that all replicates in this study were biological replicates, as defined by experiments involving different sets of animals; in no cases were the same animals tested more than once.

## Data Availability

The data that support the findings of this study are available on GitHub (https://github.com/HallemLab/Banerjee_et_al_2021).

## Acknowledgements

We thank the *Caenorhabditis* Genetics Centre, the *C. elegans* Knockout Consortium, Oliver Hobert, Cori Bargmann, Gary Ruvkun, Shai Shaham, Ikue Mori, Yuichi Iino, and Mario de Bono for strains. We thank Astra Bryant, Michelle Castelletto, and Ricardo Frausto for insightful comments on the manuscript. This work was funded by NIH F32 AI147617 (N.B.), NIH MARC T34 GM008563 (E.R.P.), NIH UF1 NS111697 (P.W.S.), and NIH R01 DC017959 and an HHMI Faculty Scholar Award (E.A.H.).

## Author Contributions

N.B., P.-Y.S., P.W.S., and E.A.H. conceived the study. N.B. and E.R.P. performed experiments.

N.B. and E.R.P. analyzed the data. N.B., E.R.P., and E.A.H. wrote the manuscript. All authors read and approved the final manuscript.

## Corresponding Author

Correspondence to Elissa A. Hallem.

---

**Fig S1.**
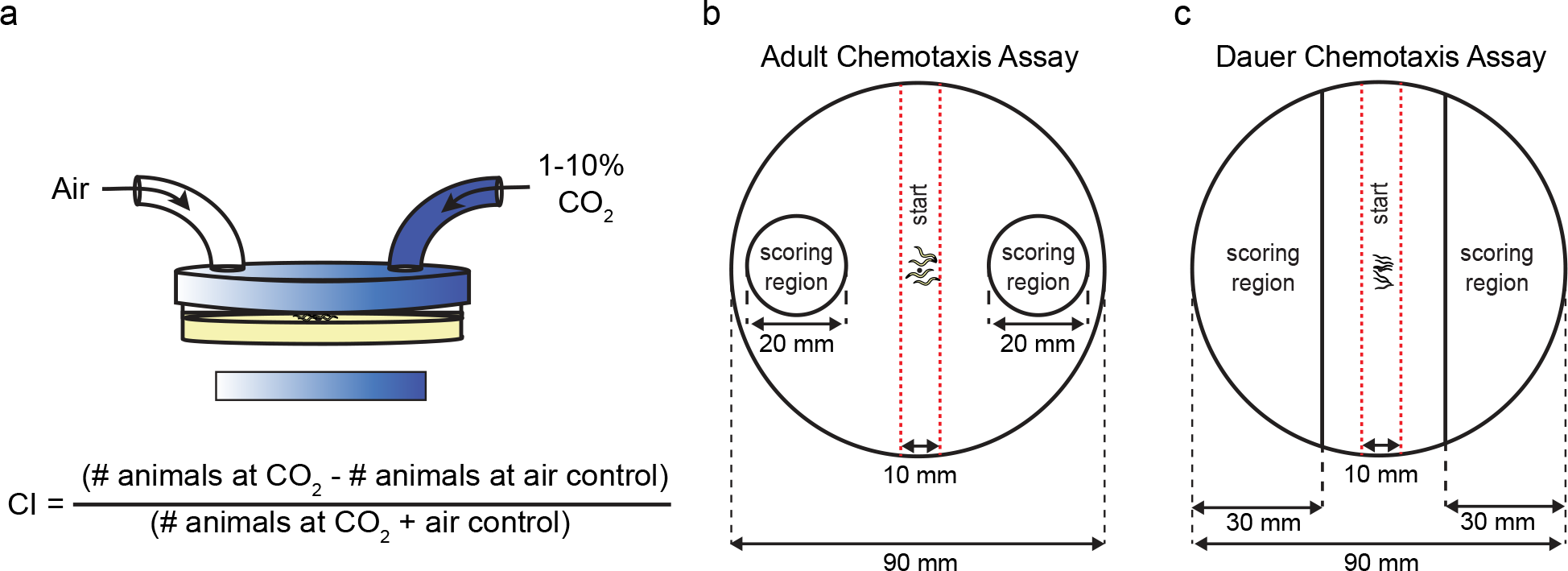
CO_2_ chemotaxis assays for *C. elegans* adults and dauers. **a.** Schematic of the CO_2_ chemotaxis assay. Animals were placed in the center of a 90 mm NGM agar plate. A CO_2_ gradient was established by delivering the specified gas mixtures through holes in either side of the plate lid. Animals were allowed to navigate in the CO_2_ gradient (indicated by the shaded rectangle below). At the end of each assay, animals on each side of the plate were counted and results were scored as a Chemotaxis Index (CI) according to the formula indicated. Adapted from Guillermin *et al.,* 2017. **b.** For adult assays, the number of animals within a 20 mm diameter circle centered under each gas inlet was counted. **c.** For dauer assays, the number of animals within the indicated 30 mm segments on both sides of the plate was counted. For b-c, dashed red lines indicate the area within which animals were placed at the beginning of the assay.

**Fig S2.**
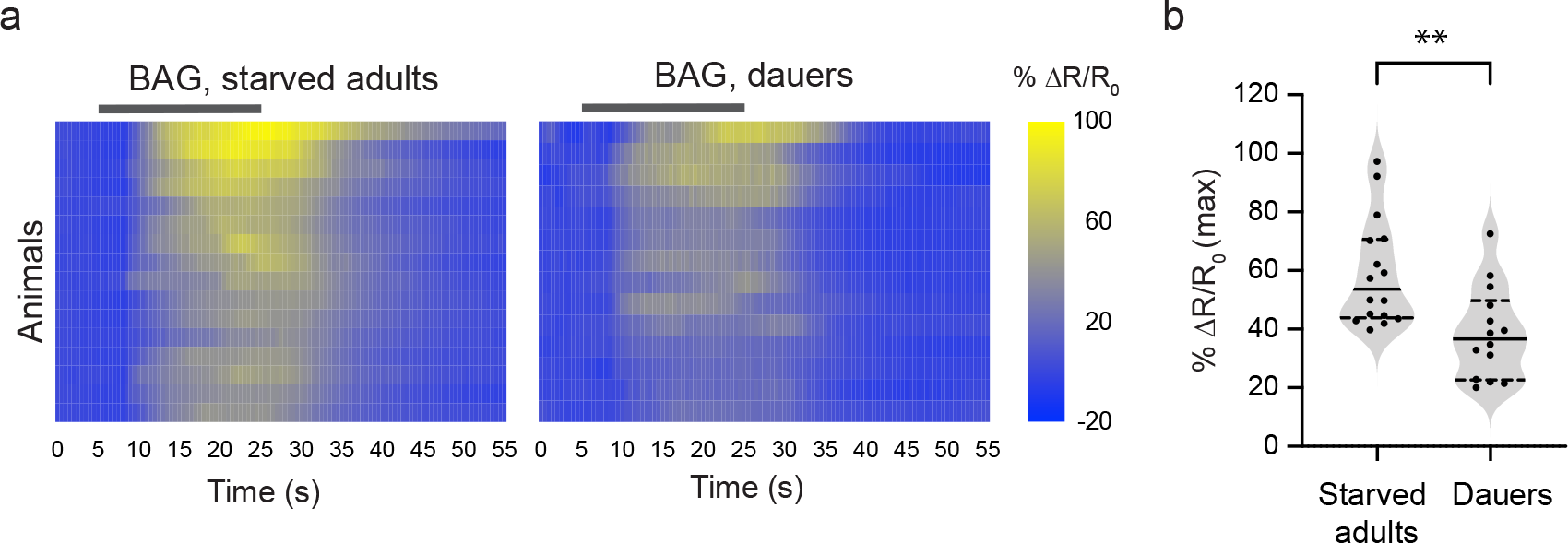
The BAG sensory neurons show similar excitatory activity in starved adults and dauers. **a.** Calcium responses in the BAG neurons of starved adults and dauers. Each row represents the response of an individual animal. Response magnitudes in the heatmaps are color-coded according to the scales (% ΔR/R_0_) shown to the right. Responses are ordered by hierarchical cluster analysis. Gray bars indicate the timing and duration of the CO_2_ pulse. **b.** Quantification of the maximum responses of BAG in starved adults and dauers. Each data point represents the response of a single animal. Solid lines in violin plots show medians and dotted lines show interquartile ranges. \**\*p*<0.01, Welch’s t-test. n = 14-16 animals per life stage. Responses are to 15% CO_2_.

**Fig S3.**
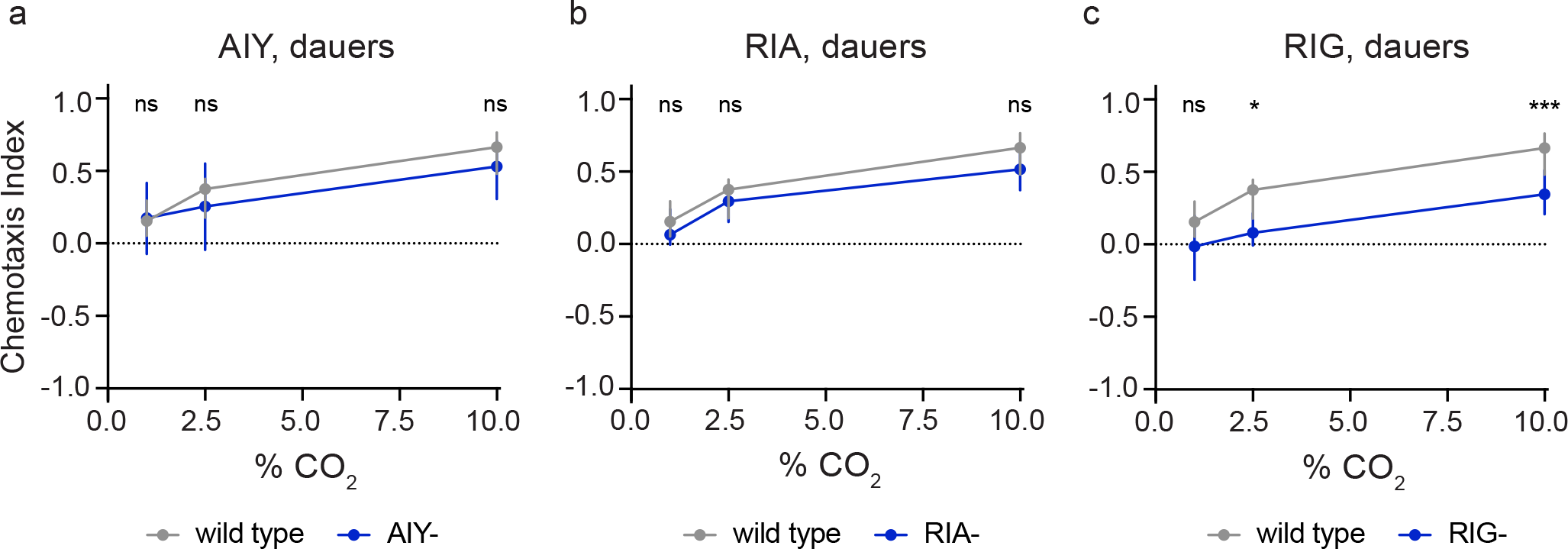
RIG interneurons, but not AIY and RIA interneurons, promote CO_2_ attraction in dauers. Behavioral responses of dauers with genetically ablated AIY (AIY-), RIA (RIA-), and RIG (RIG-) neurons to CO_2_ across concentrations. n=12-22 trials per genotype. \**p*<0.01, \*\*\**p*<0.001, ns = not significant (*p*>0.05), two-way ANOVA test with Sidak’s post-test. Graphs show medians and interquartile ranges.

**Fig S4.**
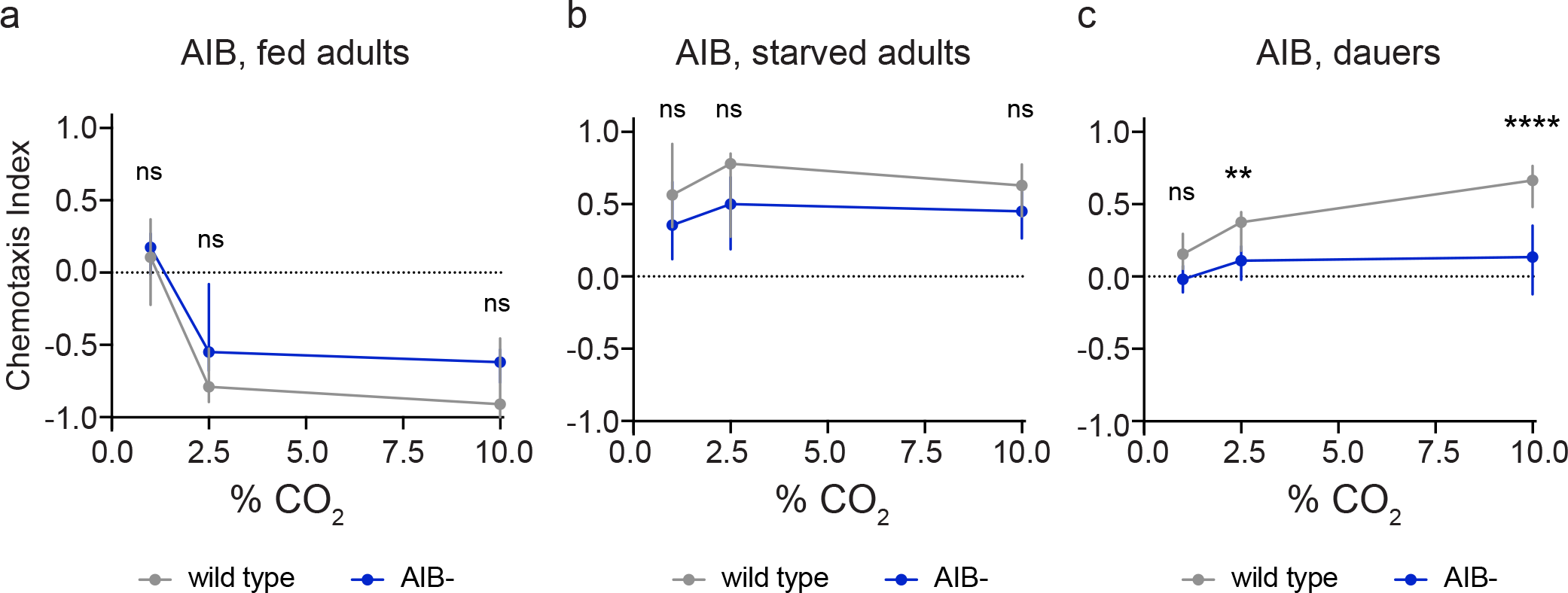
AIB interneurons promote CO_2_ attraction specifically in dauers. Behavioral responses of AIB-ablated (AIB-) well-fed adults, starved adults, and dauers in CO_2_-chemotaxis assays. \*\**p*<0.01, \*\*\*\**p*<0.0001, ns = not significant (*p*>0.05), two-way ANOVA with Sidak’s post-test. Graphs show medians and interquartile ranges. n = 12-22 trials per life stage and condition.

**Fig S5.**
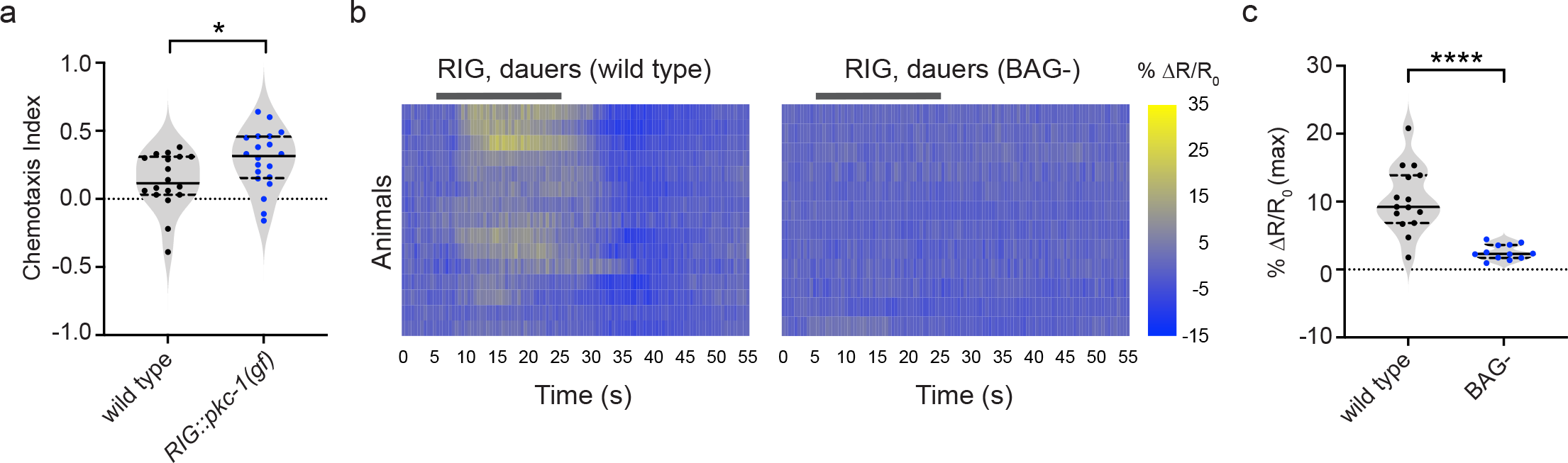
RIG activity promotes CO_2_ attraction in dauers and is dependent on BAG signaling. **a.** Dauers with RIG-specific expression of a *pkc-1(gain-of-function)* allele show significantly enhanced CO_2_ attraction compared to wild-type dauers. \**p*<0.05, Welch’s *t*-test. n = 18-20 trials per genotype. Each data point represents a single chemotaxis assay. Solid lines in violin plots show medians and dotted lines show interquartile ranges. Responses are to 1% CO_2_. **b.** Calcium responses in RIG neurons in wild-type vs BAG-ablated (BAG-) dauers. Gray bars indicate the timing and duration of the CO_2_ pulse. Each row represents the response of an individual animal. Response magnitudes in the heatmaps are color-coded according to the scale (% ΔR/R_0_) shown to the right. Responses are ordered by hierarchical cluster analysis. **c.** Quantification of the maximum responses of RIG in wild-type and BAG-dauers. Each data point represents the response of a single animal. Solid lines in violin plots show medians and dotted lines show interquartile ranges. *\*\*\*\*p*<0.0001, Welch’s *t*-test. n = 12-15 animals per genotype. For b-c, responses are to 15% CO_2_.

**Fig S6.**
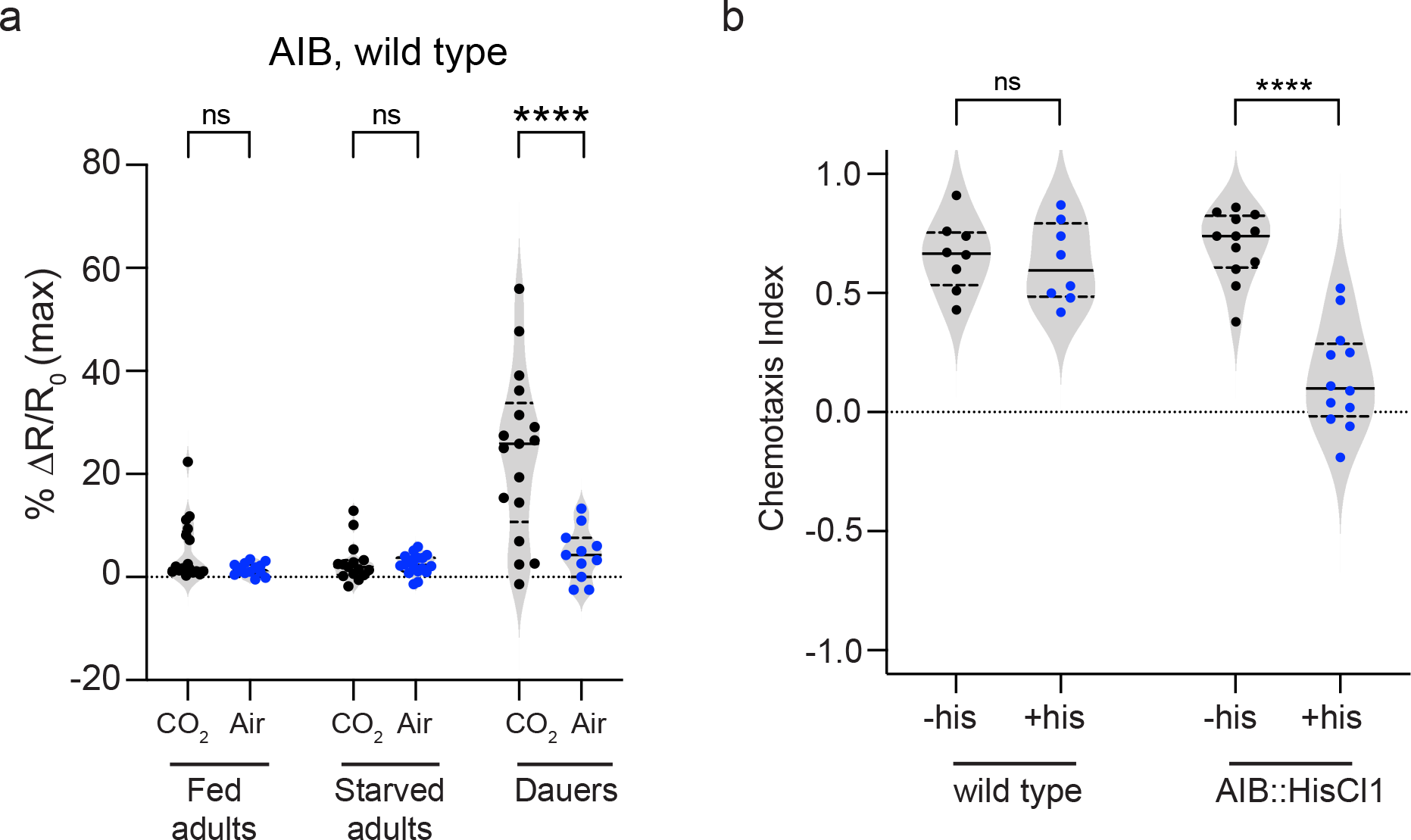
Excitatory responses in the AIB interneurons of dauers are evoked by CO_2_. **a.** Quantification of the maximum calcium responses of AIB in well-fed adults, starved adults, and dauers to CO_2_ (15% CO_2_, 21% O_2_, balance N_2_) vs. air (21% O_2_, balance N_2_). Only dauers show CO_2_-evoked activity in AIB. \*\*\*\**p*<0.0001, ns = not significant (*p*>0.5), 2-way ANOVA with Sidak’s post-test. **b.** Transient silencing of AIB neurons reduces CO_2_ attraction in dauers. Behavioral responses of wild-type dauers and transgenic dauers with AIB-specific expression of HisCl1 in CO_2_ chemotaxis assays. Dauers expressing HisCl1 in AIB (AIB::HisCl1) show significantly reduced CO_2_ attraction when treated with exogenous histamine (+his) compared to untreated controls (-his). *\*\*\*\*p*<0.0001, ns = not significant (*p* = 0.9993), two-way ANOVA with Sidak’s post-test. n = 8-12 animals per genotype and condition. Each data point represents a single chemotaxis assay. Solid lines in violin plots show medians and dotted lines show interquartile ranges. Responses are to 10% CO_2_.

**Fig S7.**
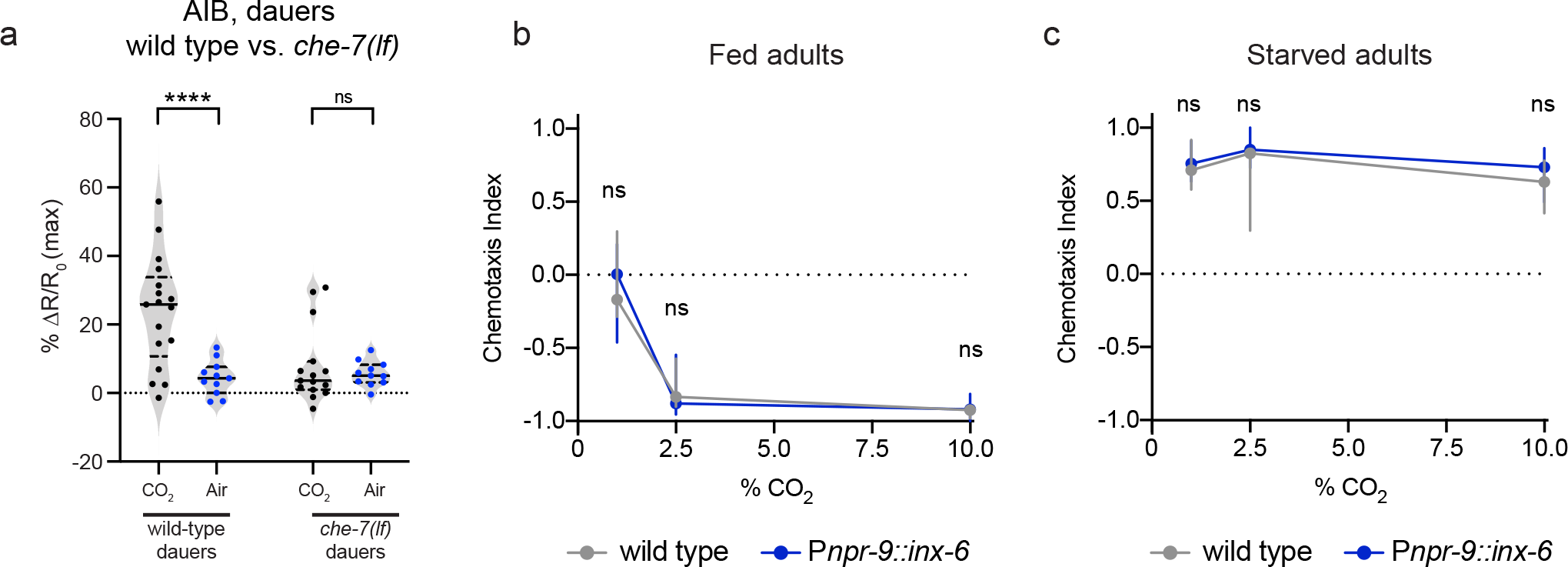
The BAG-AIB gap junction is required for CO_2_-evoked activity in AIB and modulates CO_2_-evoked behavior specifically in dauers. **a.** Quantification of the maximum calcium responses of AIB in *che-7(ok2373)* mutants in response to CO_2_ vs. air. Wild-type but not *che-7(lf)* dauers show CO_2_-evoked activity in AIB. *\*\*\*\*p*<0.0001, ns = not significant (*p* = 0.8571), two-way ANOVA with Sidak’s post-test. For a-b, each data point represents the response of a single animal. Solid lines in violin plots show medians and dotted lines show interquartile ranges. n = 11-18 animals per genotype and condition. **b-c.** Ectopic expression of INX-6 in AIB does not alter CO_2_ responses in adults. Behavioral responses of well-fed adults (**b**) and starved adults (**c**) expressing *inx-6* specifically in AIB under the control of the *npr-9* promoter (P*npr-9*::*inx-6*) in CO_2_-chemotaxis assays across concentrations. n = 10-16 animals per genotype and condition. Lines show medians and interquartile ranges. ns = not significant (*p*>0.1), two-way ANOVA with Sidak’s post-test.

**Fig S8.**
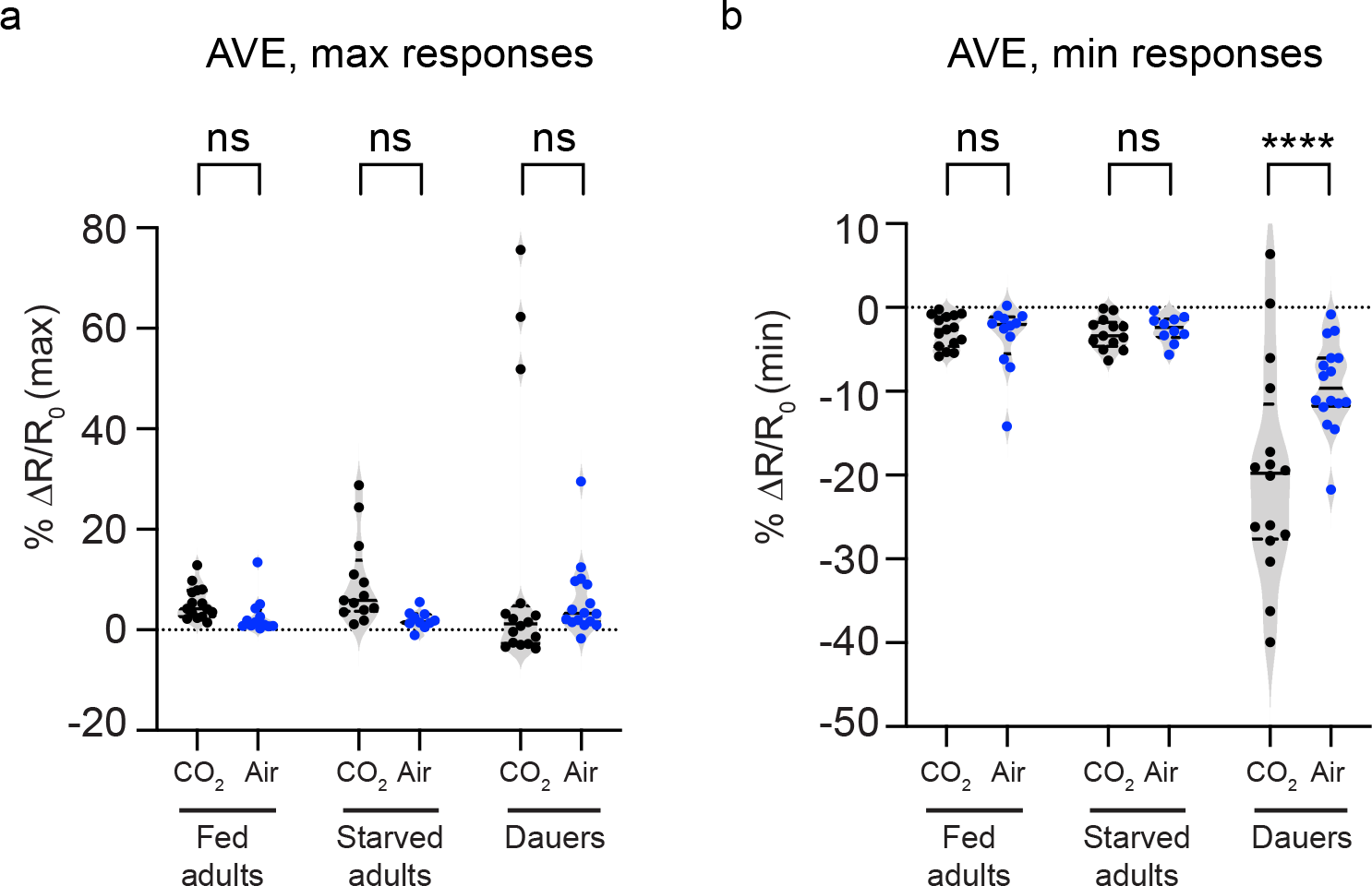
Inhibitory responses in the AVE interneurons of dauers are evoked by CO_2_. Quantification of the maximum (**a**) and minimum (**b**) calcium responses of AVE in well-fed adults, starved adults, and dauers to CO_2_ (15% CO_2_, 21% O_2_, balance N_2_) vs. air (21% O_2_, balance N_2_). Only dauers show CO_2_-evoked inhibitory activity in AVE. In a-b, each data point represents the response of a single animal. Solid lines in violin plots show medians and dotted lines show interquartile ranges. *\*\*\*\*p*<0.0001, ns = not significant (*p*>0.4), two-way ANOVA with Sidak’s post-test. n = 11-16 animals per life stage and condition.

**Fig S9.**
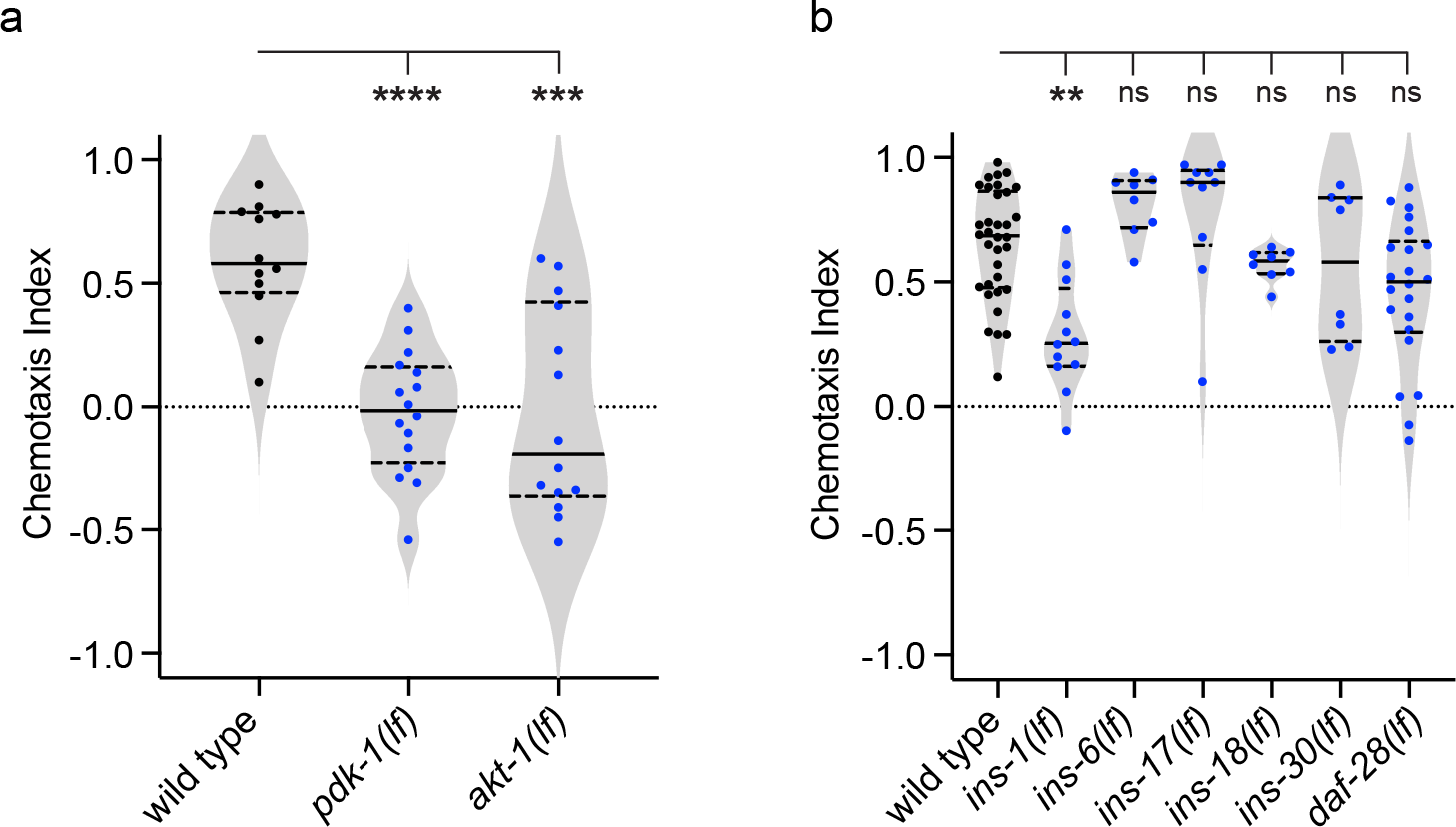
Insulin signaling promotes CO_2_ attraction in dauers. **a.** Behavioral responses of *pdk-1(sa709)* and *akt-1(mg306)* mutant dauers in CO_2_ chemotaxis assays. *\*\*\*\*p*<0.0001, *\*\*\*p*<0.001, one-way ANOVA with Dunnett’s post-test. n = 12-16 animals per genotype. **b.** The neuropeptide INS-1 promotes CO_2_ attraction in dauers. Behavioral responses of dauers carrying loss-of-function mutations in candidate insulin-like peptide genes in CO_2_ chemotaxis assays. \*\**p*<0.01, ns = not significant (*p*>0.2), Kruskal-Wallis test with Dunn’s post-test. n = 8-34 animals per genotype. The specific loss-of-function (*lf*) alleles used for individual insulin-like peptide genes are indicated in the Materials and Methods. For a-b, each data point represents a single chemotaxis assay. Solid lines in violin plots show medians and dotted lines show interquartile ranges. Responses are to 10% CO_2_.

**Fig S10.**
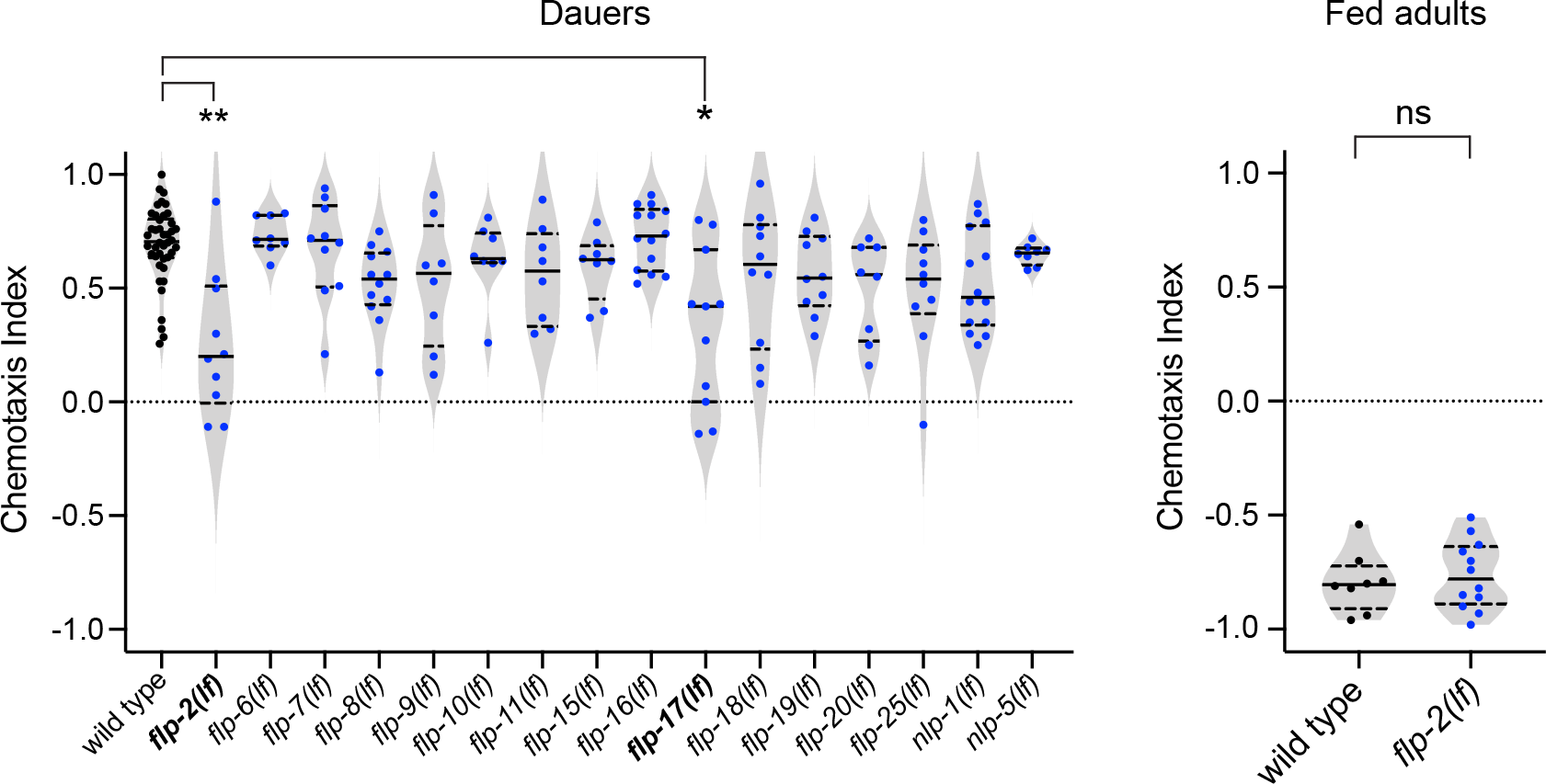
A reverse genetic screen for neuropeptides that regulate CO_2_ attraction in dauers. **a.** FLP-2 and FLP-17 neuropeptides regulate CO_2_ attraction in dauers. The graph shows the behavioral responses of dauers with loss-of-function mutations in candidate neuropeptide genes in CO_2_ chemotaxis assays. \*\**p*<0.01, \**p*<0.05, Kruskal-Wallis test with Dunn’s post-test. n = 8-42 assays per genotype. The specific loss-of-function (*lf*) alleles used for individual neuropeptide genes are indicated in the Methods. **b.** FLP-2 neuropeptides do not regulate CO_2_ repulsion in well-fed adults. Behavioral responses of *flp-2(gk1039)* mutant well-fed adults in a CO_2_ chemotaxis assay. ns = not significant (*p* = 0.6176), Welch’s *t*-test. n = 8-12 assays per genotype. For a-b, each data point represents a single chemotaxis assay. Solid lines in violin plots show medians and dotted lines show interquartile ranges. Responses are to 10% CO_2_.

**Table S1.**
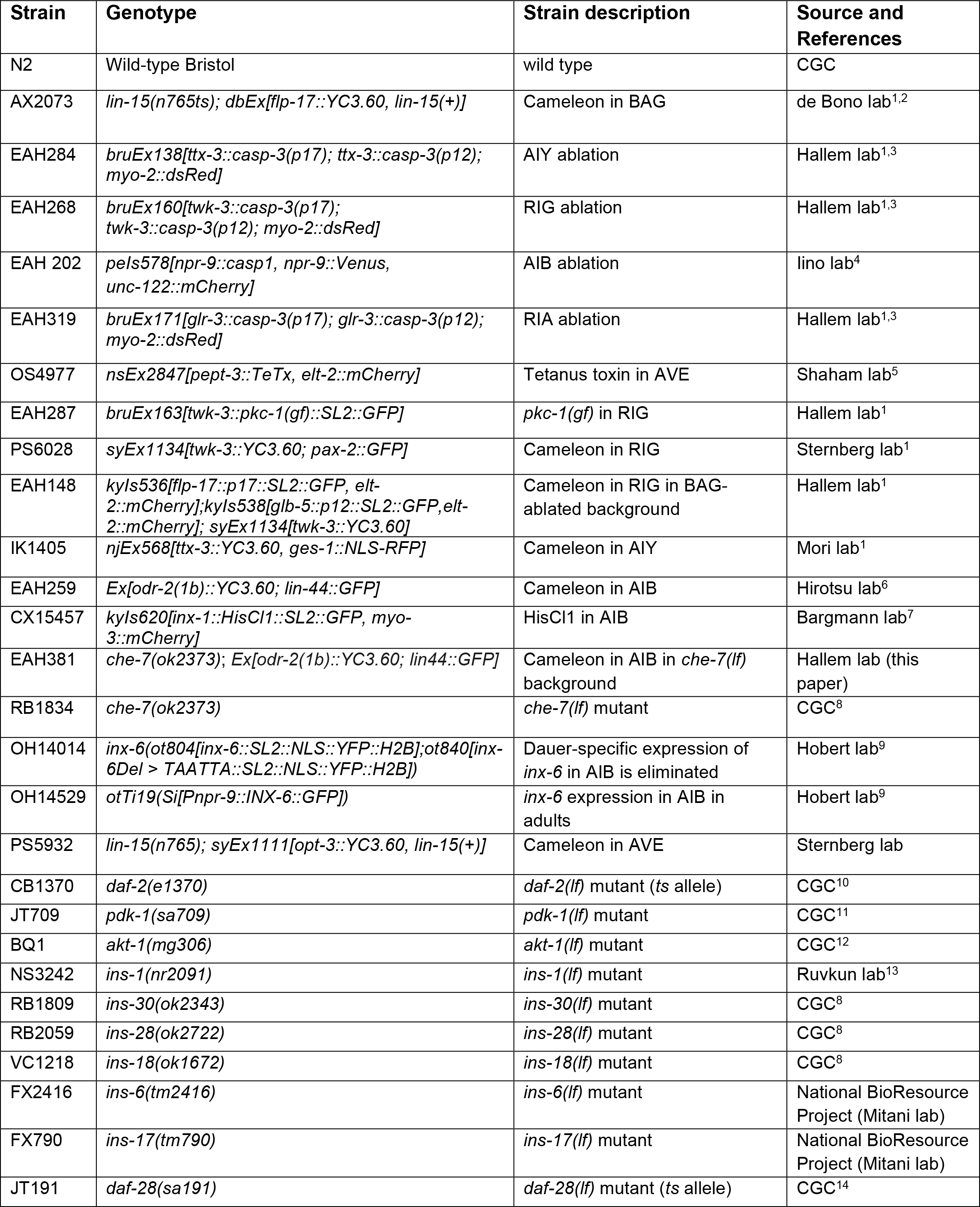
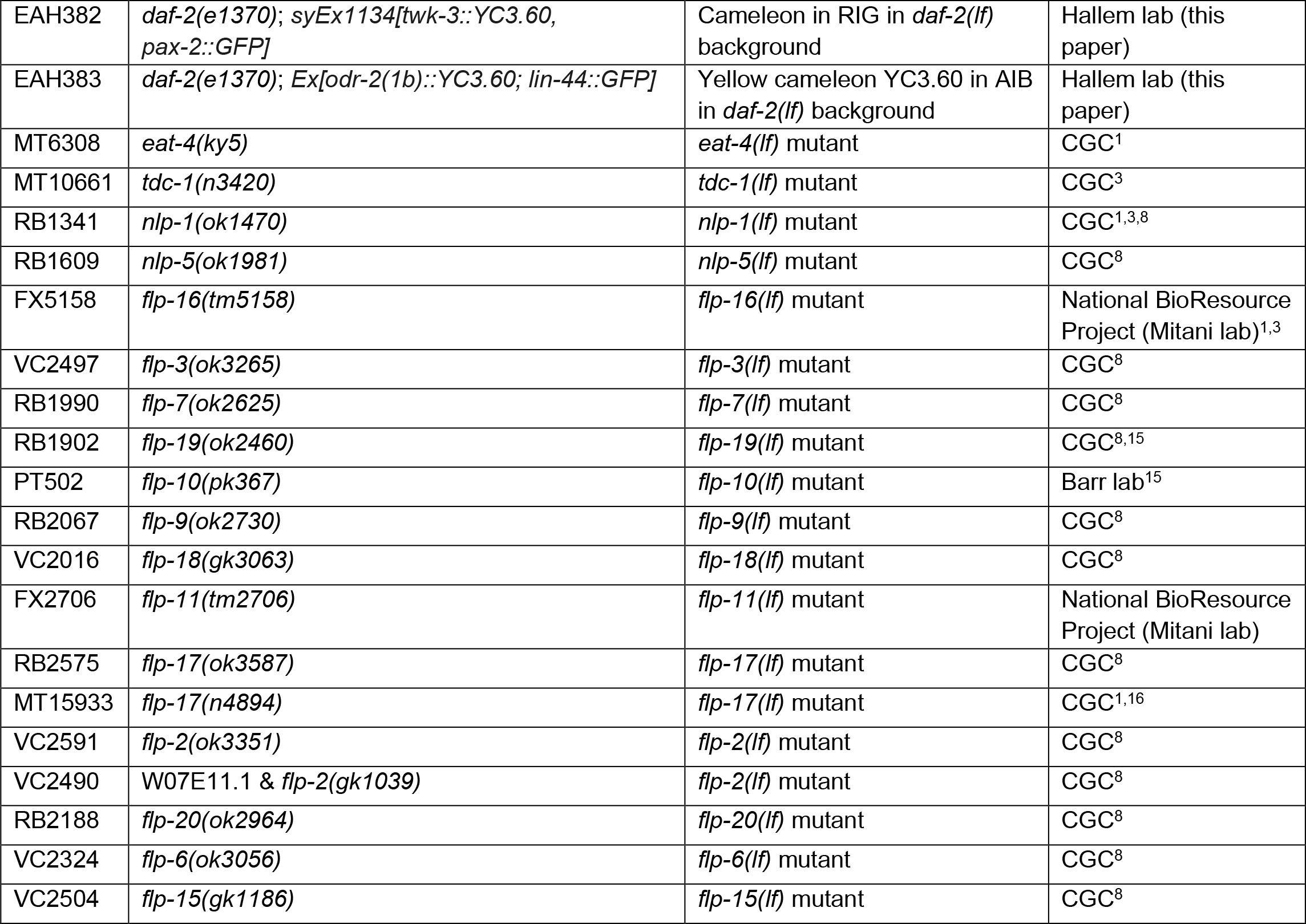
The list of strains used. *lf* = loss-of-function mutation; *ts* = temperature-sensitive.

